# Selectivity for high-level language processing is highly localized in individual brains

**DOI:** 10.64898/2026.07.16.738967

**Authors:** Rebecca M. Belisle, Terri L. Scott, Tyler K. Perrachione

## Abstract

Some aspects of human behavior and cognition depend on focal and selective cortical areas, such as the frontal eye fields or fusiform face area, while others, like semantic knowledge, are broadly distributed across the cortex. Whether higher-level cognitive functions like language can also be highly localized has been a longstanding matter of debate. Here, we provide multiple lines of evidence that receptive language in the brain is subserved by a network of discrete, focal, and uniquely language-selective areas when examining individual brains. Using precision neuroimaging, we observed highly circumscribed patches of cortex that are distinctly selective for language, uniquely consistent in their response properties during language processing, and highly reliable in their anatomical locations within individuals (though variable in location across individuals). These findings indicate that language regions in the brain are characterized by unique functional profiles and sharp boundaries, consistent with the systems neuroscience definition of true “cortical areas.”

## Introduction

Two competing models have inspired and shaped efforts to understand the functional organization of the human brain (Dick et al., 2004). One longstanding view holds that the brain has a highly modular organization with distinct cortical areas subserving specific mental functions, like perceiving faces (Kanwisher et al., 1997; Levakov et al., 2022; Petersen et al., 2024), controlling eye movements (Bedini et al., 2023; Lobel et al., 2001), and, putatively, processing language (Chai et al., 2016; Fedorenko et al., 2024; Haglund et al., 1994; Ojemann et al., 1989). Functional modularity of the brain’s cerebral cortex parallels neuroanatomical observations of microstructural arealization, from Brodmann’s well-known parcellation (Brodmann, 1909) to modern cytoarchitectural approaches (Amunts et al., 2007; Bruno et al., 2022). The alternative view holds that, the brain is organized along functional gradients, such that, instead of sharp functional boundaries, tissue supporting one operation blends gradually into the tissue supporting other operations (Bernhardt et al., 2022; Dohmatob et al., 2021; Huntenburg et al., 2018) or that cognitive faculties are distributed broadly across the brain (Drijvers et al., 2025; Lashley, 1929; Noble et al., 2024; Westlin et al., 2023). This debate parallels a classic question from cognitive science regarding the modularity of mental operations (Fodor, 1983; Carruthers, 2006; Prinz, 2006).

The human capacity for language is among the most widely discussed in terms of its modular vs. nonmodular organization, both with respect to its underlying mental operations (Drakoulaki et al., 2024; Federmeier et al., 2020; Glucksberg et al., 1973; Petkov et al., 2020; Yang & Piantadosi, 2022) and their neurobiological implementation (Fedorenko & Shain, 2021; Fedorenko & Varley, 2016; Fedorenko et al., 2024; Klimovich-Gray & Bozic, 2019; Price et al., 2005). The neural modularity of language is also one of the earliest debates in human neuroscience, with lesion-based studies like those of Broca and Wernicke suggesting that specific brain regions were essential to particular facets of language (Broca, 1861; Broca, 1865; Wernicke, 1874), while others argued against strict localization of cognitive functions (Goltz, 1888; Müller, 1837; Lashley, 1929). Subsequent work using electrical stimulation during brain surgeries also supported the functional localization of language: Stimulation of focal brain areas impaired object naming, but stimulation of immediately adjacent cortex did not impact expressive language behaviors (Ojemann et al., 1989) and often caused responses in non-linguistic domains (e.g., specific somatic sensations; Penfield & Jasper, 1954). Notably, the exact locations of these anomia-inducing stimulation sites in the frontal and temporal lobes varied across patients, but resecting tissue at least one centimeter away from such a site preserved language abilities after surgery (Haglund et al., 1994; Ojemann et al., 1989).

Further evidence for the existence of a network of focal areas specializing in language comes from recent single-subject analyses of fMRI data, in which functional regions of interest are defined independently in individual brains and then characterized by their responses to various tasks and cognitive domains (Devaney et al., 2015; Fedorenko & Blank, 2020; Henderson et al., 2022; Saxe et al., 2006; Wolna et al., 2024; 2026). There is now a wealth of evidence indicating that, when functionally defined in individual brains, areas can be found that are uniquely selective for language as opposed to tasks involving math (Fedorenko et al., 2011), spatial working memory (Fedorenko et al., 2011; Lee et al., 2024), theory of mind (Shain et al., 2023), object categorization (Benn et al., 2023), intuitive physical reasoning (Kean et al., 2025), domain-general processing (Fedorenko et al., 2012), music (Chen et al., 2023), and numerous other domains (Ivanova et al., 2020; Pritchett et al., 2018). While these language areas’ functional profiles are similar across individuals, and they lie within the same broad anatomical regions like the inferior frontal or lateral temporal lobes, their precise locations with respect to the cortical curvature vary enormously from person-to-person (Lipkin et al., 2022). Further evidence that these focal areas comprise a coherent “language network” comes from their correlated resting-state fMRI activity patterns (Braga et al., 2020; Kong et al., 2019), with the areas in this task-free language network also showing functional selectivity for language when interrogated using task fMRI (Du et al., 2024; Salvo et al., 2025). In contrast to historical lesion-based studies or classical group-average fMRI findings, where both linguistic and non-linguistic faculties are attributed to broad anatomical regions like the IFG (Blumstein, 2009; Price, 2012; Scott, 2019), contemporary findings are beginning to provide a more precise and individualized picture of language in the brain, in which there appears to be a network of discrete regions that are functionally specialized for language (Fedorenko & Blank, 2020; Fedorenko et al., 2024).

However, prior work showing that some areas are strictly language-selective does not, in and of itself, refute the possibility that there are gradients of language selectivity (i.e., a gradual reduction in language selectivity centered on these focal peaks), nor do localized peaks preclude the possibility that there might be some areas that are moderately engaged by both language and other domains. By investigating only the most (or least) language-selective voxels, prior fMRI studies of focal language localization leave unanswered the fundamental questions of whether and to what extent these language nodes are functionally distinct from immediately adjacent tissue, and therefore to what extent language is organized in a modular vs. gradient fashion across the entire brain.

Here, we test these competing models of localized vs. gradient functional organization of language across large swaths of the cerebral cortex (**Fig. 1**). In particular, we examined whether cortical selectivity for language in individual brains is (i) *gradiently organized*, such that functional selectivity for language decreases more or less linearly with greater distance from local peaks, suggesting that there are no clear boundaries distinguishing language-selective brain tissue from tissue responsive to other cognitive domains; (ii) *widely distributed* across large swaths of cortex, suggesting that core language selectivity is more extensive than only the highly localized areas examined in prior fMRI studies, or (iii) *distinctly heightened in focal areas*, suggesting a predominately modular organization of core language-selective areas. To test these hypotheses, we employed a naturalistic language-listening fMRI paradigm that reliably identifies regions that are strongly and specifically selective for language in individual brains based on the contrast of intact vs. acoustically degraded, unintelligible speech (Scott et al., 2017; Lee et al., 2024). Instead of using an arbitrary activation threshold to determine which voxels to include in functional regions of interest (fROIs), we looked beyond the most language-selective cortex to investigate how language activation varied continuously over wide brain areas, as well as how contiguous and coherent (i.e., how “area-like”) these patterns of language-selective activation were. To test whether language regions formally adhere to the functional criteria used to define “cortical areas” in systems neuroscience (Cadwell et al., 2019; Eickhoff et al., 2018; O’Leary et al., 2007; Passingham et al., 2002; Petersen et al., 2024; Rakic, 1988), we examined the *focality* of language activation in the brain, as well as these areas’ test-retest *reliability*, *spatial consistency*, and *spatial contiguity* (i.e., connectedness) of voxels with similar levels of language selectivity. Ultimately, our findings support the view that high-level receptive language processing in the brain enlists a network of selectively responsive, highly focal areas that are situated uniquely with respect to the cortical curvature in individual brains.

**Figure 1:**
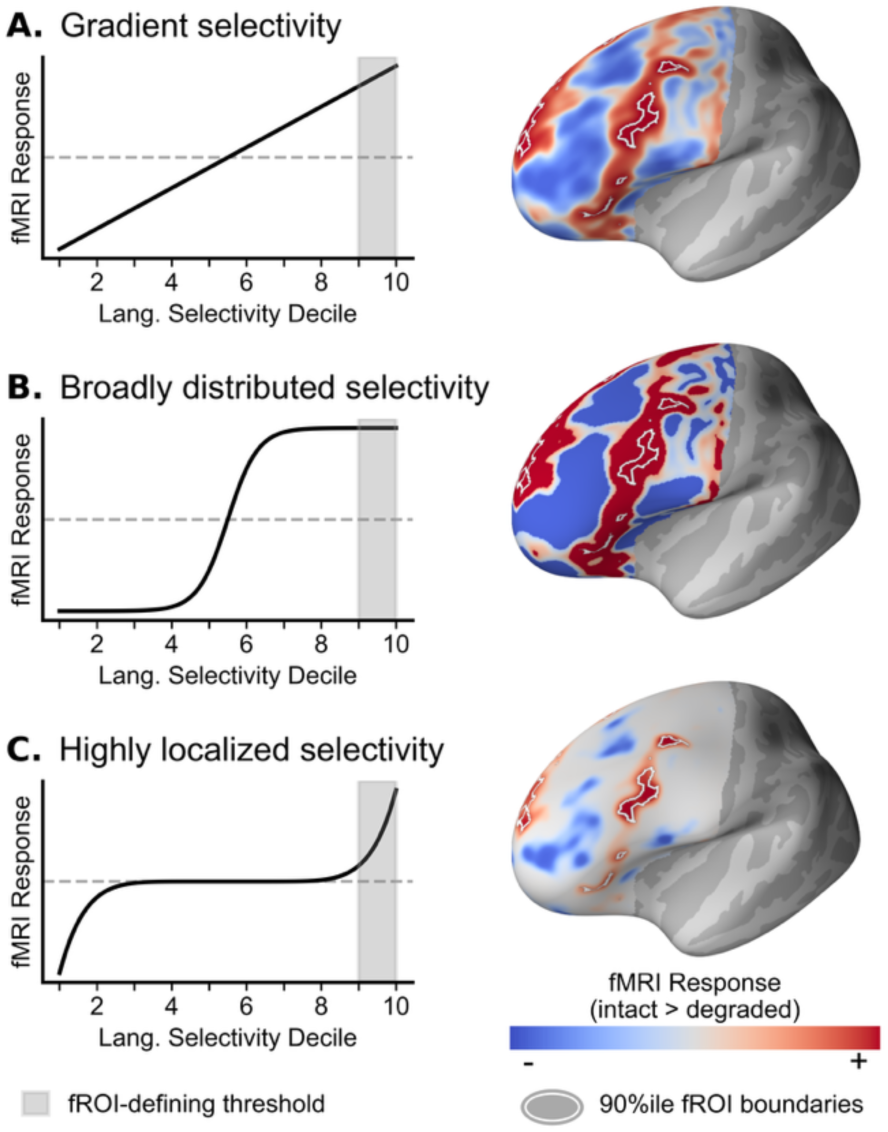
Hypothetical language response profiles of fMRI activation in left frontal lobe as a function of language selectivity rank (decile). **(A)** Gradient selectivity, with the language response profile appearing mostly linear, reflecting varying degrees of language response across the cortex. **(B)** Distributed selectivity, with a large proportion of cortex strongly responsive to language. **(C)** Focal selectivity, with strong language responses concentrated in a relatively small portion of cortex, reflected in response curves mostly near zero but with a steep peak at the most-selective end of the distribution.

## Results

### Task activation

We observed unique patterns of activation not only across tasks, but also across subjects. In the frontal lobe (**Fig. 2A**), the strongest activation to language was typically found concentrated along inferior frontal gyrus (IFG), with additional activation prominent in anterior superior frontal gyrus (SFG) and posterior middle frontal gyrus (MFG) into precentral gyrus (PrCG) (**Extended Data Fig. 1A**). In contrast, the strongest activation to spatial and verbal working memory tended to be more widely distributed throughout ventral IFG, anterolateral MFG, and dorsal SFG (**Extended Data Fig. 1B,C**). Within subjects, regions most responsive to language appeared to be distinct from those responsive to the working memory tasks, which in turn recruited many regions in common. In the temporal lobe (**Fig. 3A**), the strongest activation for language was typically found in anterior STG and along mid-to-posterior STS (**Extended Data Fig. 1A**). Activation for verbal working memory was often around transverse temporal gyrus and posterior STG (**Extended Data Fig. 1B**). For the spatial working memory contrast, activation spanned parts of posterior MTG and ITG (**Extended Data Fig. 1C**). Finally, in the parietal lobe (**Fig. 4A**) the strongest language activation was typically concentrated near the temporoparietal junction in ventral supramarginal gyrus (SMG) and anterior inferior parietal lobule (IPL). Activation to working memory tasks was more prominent throughout parietal lobe, with verbal working memory response concentrated most strongly in SMG and the strongest spatial working memory responses in superior parietal regions (**Extended Data Fig. 1C**). Broadly, these findings reflect the established knowledge about the cortical organization of these tasks; therefore, in order to ascertain the focality or overlap of these tasks in the same brain tissue, we also quantitatively investigated the spatial organization of these response profiles in individual brains.

**Figure 2:**
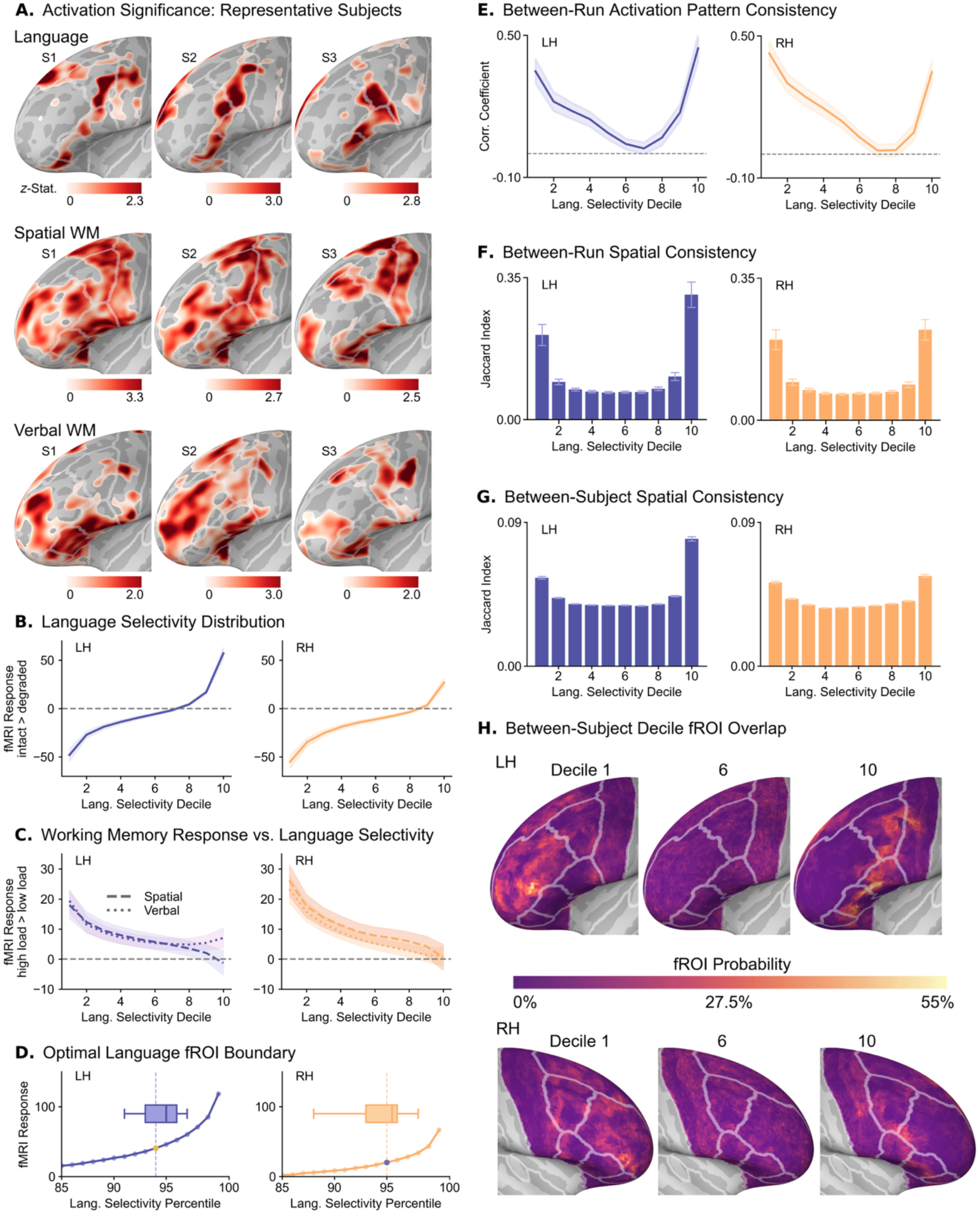
Frontal lobe. **(A)** Left hemisphere fMRI task activation significance (GLM z-statistic) maps for three example individuals with Desikan-Killiany atlas regions outlined. **(B)** The average fMRI response magnitude (contrast parameter estimate) as a function of language selectivity (grouped by decile) for the language localizer, as well as for the **(C)** spatial working memory and verbal working memory tasks. **(D)** Average fMRI response magnitude to language, focusing on only the most selective voxels (grouped by percentile), showing the location of the sharp change (“knee point”) in language selectivity at the group average level (average of individuals’ fMRI response values for each percentile; dashed lined indicating knee at: LH: 93%ile, RH: 94%ile) and at the individual level (box plot of individuals’ knee values; median: LH: 94%ile, RH: 94.5%ile). **(E)** The correlation between the voxelwise language activation patterns within language-selectivity deciles across runs. **(F)** Quantified test-retest within-subject spatial overlap for language-selectivity deciles across runs. **(G)** Quantified spatial overlap for language-selectivity deciles across different subjects in standard space. **(H)** Probability maps of the 1^st^, 6^th^, and 10^th^ deciles across subjects. The value indicates the proportion of subjects whose decile fROI includes that vertex. Two runs of fROIs were included per subject. **(B-C, E-G)** Shading and error bars represent the standard error of the mean across subjects.

**Figure 3:**
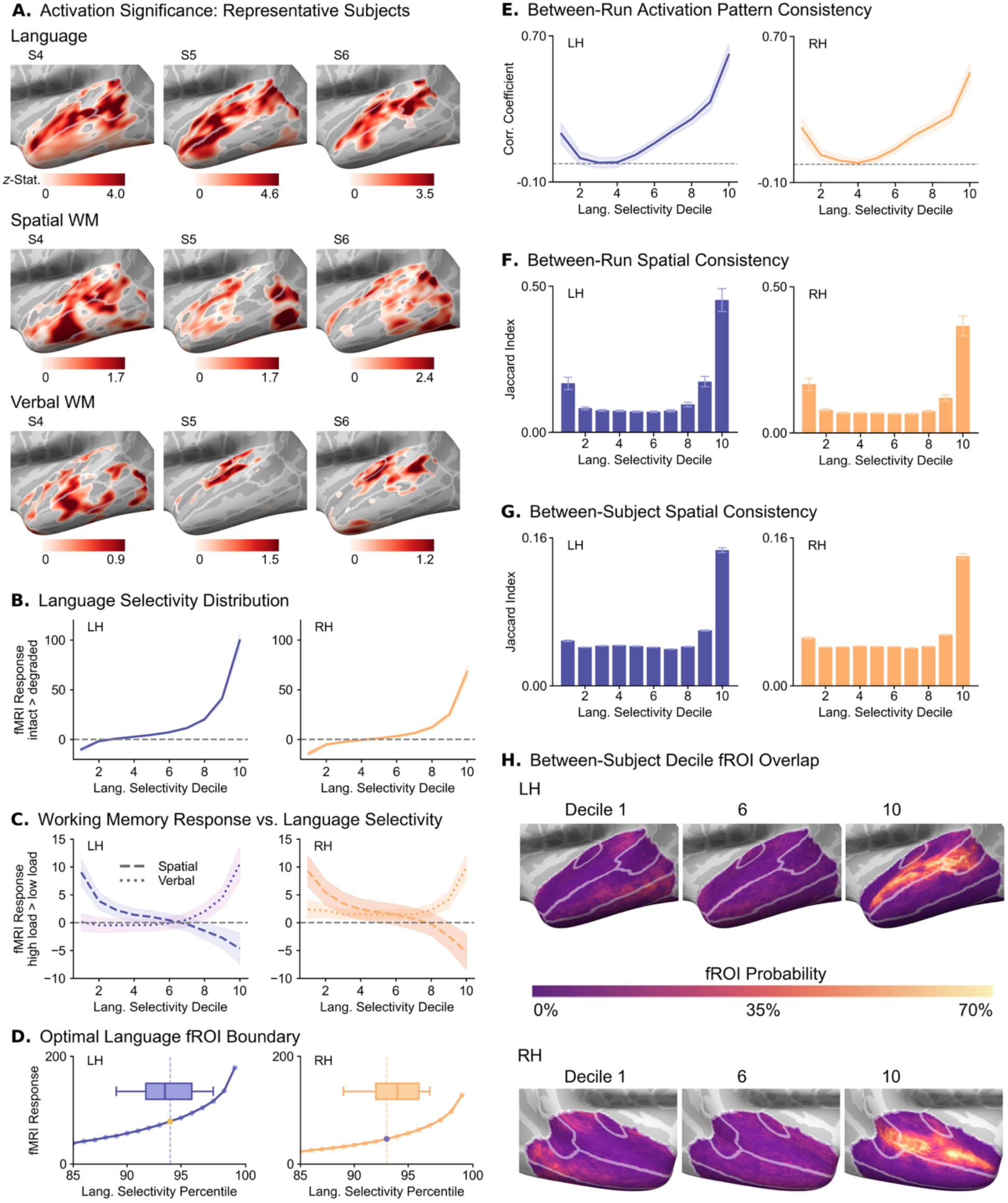
Temporal lobe. **(A – C, E – H)** Same descriptions as in Figure 2. **(D)** The knee point for language selectivity at the group average level (average of individuals’ fMRI response values for each percentile; dashed lined indicating knee at: LH: 93%ile, RH: 92%ile) and at the individual level (box plot of individuals’ knee values; median: LH: 92.5%ile, RH: 93%ile).

**Figure 4:**
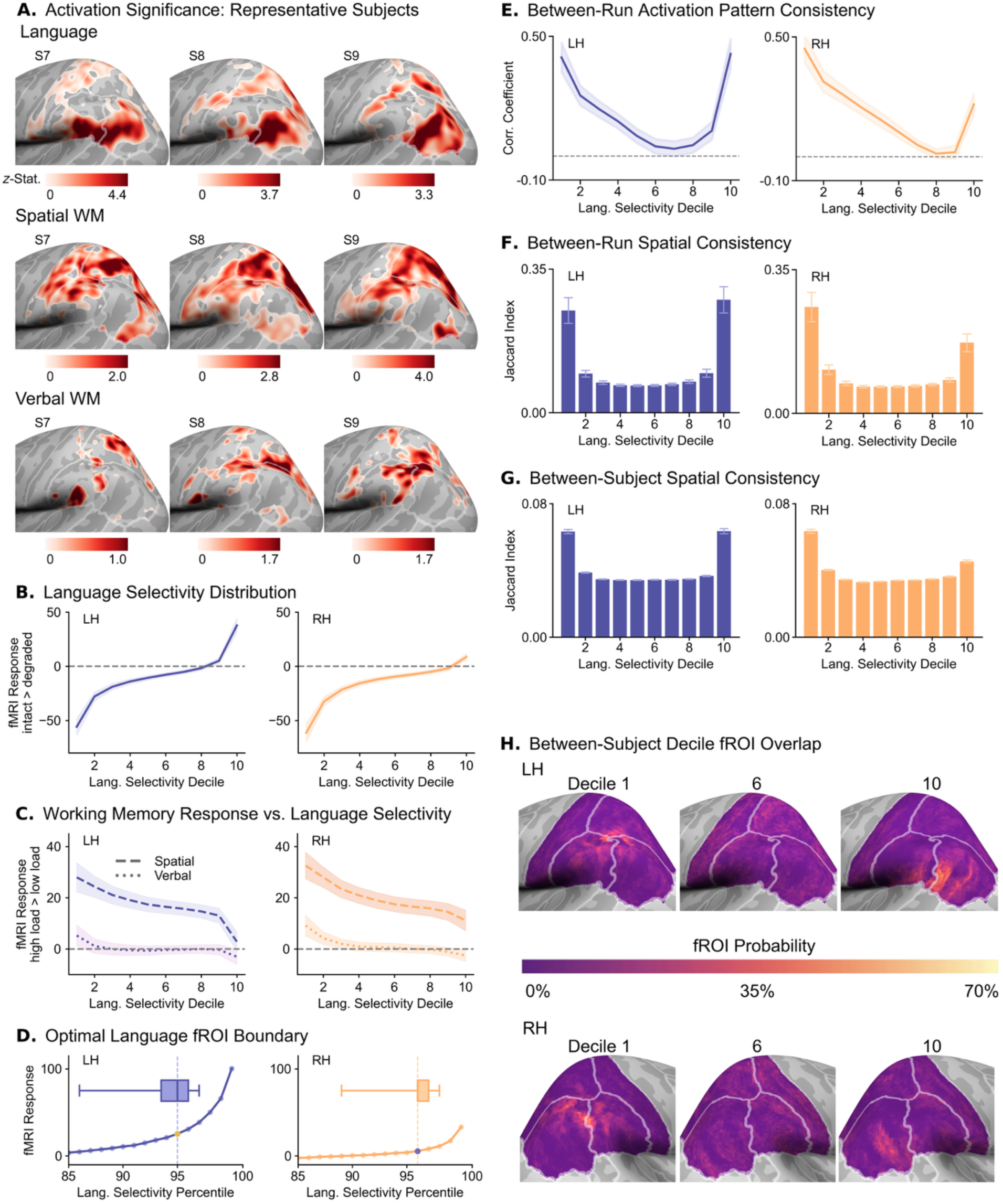
Parietal lobe. **(A – C, E – H)** Same descriptions as in Figure 2. **(D)** The knee point for language selectivity at the group average level (average of individuals’ fMRI response values for each percentile; dashed lined indicating knee at: LH: 94%ile, RH: 95%ile) and at the individual level (box plot of individuals’ knee values; median: LH: 94%ile, RH: 95%ile).

### Localized peaks of highly language-selective fMRI response

To determine whether language selectivity follows gradient, distributed, or localized cerebral organization (**Fig. 1**), we examined the fMRI response magnitude of the language localizer contrast (listening to intact > degraded speech) for all cortical voxels in the frontal (**Fig. 2B**), temporal (**Fig. 3B**) and parietal (**Fig. 4B**) lobes as a function of their ranked response to that task (by decile). We found that the top language-selective voxels (90%ile) in each of the three lobes exhibited a distinctly heightened language response compared to the next-most-selective voxels (80%ile), with the lowest-responsive voxels (0%ile) exhibiting a distinctly negative peak. The cortical language response profile was distinctly nonlinear, with a significantly cubic relationship as a function of activation rank across all frontal, temporal, and parietal regions examined (*β* > 0, *p* < 0.001; **Supplementary Table 1**). This motif of focally increased responses was also conserved when examining the entire left-hemisphere cerebral cortex at once (**Extended Data Fig. 2**), as well as in more circumscribed atlas regions (**Supplementary Figs. 1B–6B**). In the right hemisphere, the shapes of the language response profiles were similar to those in the left, but with smaller absolute response magnitudes among the top decile fROIs. This pattern of results is consistent with the hypothesis of focally localized language-selective brain tissue (**Fig. 1C**) and does not reflect the alternative hypothesized organizations of either gradient selectivity or broad swaths of language-selective cortex. Critically, this finding of a small number of highly language-selective voxels was observed in each individual brain (**Extended Data Fig. 3**), indicating that localized selectivity is not an artifact of averaging across subjects, but a common organizing principle of language in the brain.

### Only the most language-selective areas are not responsive to other tasks

We also examined how responses to spatial and verbal WM (contrasting high > low working memory load) varied as a function of language selectivity. Across the lobes, we found that spatial WM response profiles tended to reverse the language response profiles: The most language-selective regions were either negatively or non-responsive to the spatial WM task, with spatial WM response increasingly gradually with decreasing language response (**Fig. 2C-4C**).

To ensure that the shapes of the language-response curves (**Fig. 2B-4B**) were unique to the language localizer contrast (and not characteristic of a region regardless of task), we also analyzed the fMRI response magnitude vs. task-selectivity rank for the spatial WM task (**Supplementary Fig. 7**). While spatial WM response magnitudes were also heightened among the most selective voxels, this nonlinearity was much less than what we saw for language, with a more graded pattern of responses among the intermediately SWM-selective voxels compared to the near-zero responses seen in the language-response curves.

Interestingly, the relationship between language selectivity and verbal WM varied by lobe: In left and right frontal lobes, verbal WM activation was strongest in the least language-selective areas and decreased gradually with increasing language selectivity (**Fig. 2C**). However, in left frontal lobe, the most language-selective voxels also showed a slight increase in verbal WM response. Verbal WM responses in bilateral temporal lobes were notably different from the inverse relationship with language selectivity found elsewhere, with the top language-selective regions also showing the strongest response to verbal WM load, and little verbal WM response in regions with weak language response (**Fig. 3C**). The relatively weak verbal WM response in the parietal lobe was concentrated in the least language-selective areas (**Fig. 4C**).

### Individual selectivity thresholds

At first glance, the dramatic decrease in language selectivity from voxels in the 90%ile to those in the 80%ile deciles seems to affirm the widespread use of “top 10%” as the fROI-forming threshold in prior group-constrained subject-specific (GCSS) precision neuroimaging studies of language (e.g., Chai et al., 2016; Kunert et al., 2015; Malik-Moraleda et al., 2022; Ozernov-Palchik et al., 2026; Scott et al., 2017; Van Kube et al., 2025; Wolna et al., 2024). However, this *a priori* threshold potentially obscures differences in the focality of language-selectivity across regions or individuals. We used a data-driven thresholding approach to identify the boundary of language-selective voxels for each individual, motivated by prior findings of individual differences in relative cortical area size (Dworetsky et al., 2024; Glasser et al., 2016). We examined voxelwise response properties more granularly, here using language-selectivity *percentile* (instead of decile) and, for each subject, identifying the point of maximum curvature where language response starts increasing more rapidly (**Fig. 2D–4D**). Individual language-selectivity forming thresholds ranged from 90-96%ile (mean=93.4%ile) in left frontal lobe, 87-97%ile (mean=93.6%ile) in right frontal lobe, 88-97%ile (mean=92.6%ile) in left temporal lobe, 88-96%ile (mean=92.6%ile) in right temporal lobe, 85-96%ile (mean=93.4%ile) in left parietal lobe, and 88-97%ile (mean=94.7%ile) in right parietal lobe.

In precision neuroimaging and GCSS studies of language, the search space for individual subjects’ fROIs is often restricted to within group- or population-defined functional “parcels” that capture the probabilistic extent of activation across many subjects (Lipkin et al., 2022). We examined how such prior constraint to areas most likely to have language-related activation affected the focality of language responses compared to those measured across the entire lobe. We created 13 group-constrained functional parcels, delineating cortical surface vertices where significant language localizer activation was likely to be found across all subjects in our sample, and repeated the data-driven subject-specific fROI-forming threshold analysis within these regions (**Extended Data Fig. 4**). Here, the knee points in the language-response curves were also evident, with subject-specific fROI thresholds (71-97%ile) tending to be more inclusive in key frontotemporal functional parcels than those based on anatomical atlas-defined regions, suggesting that *a priori* fROI thresholding at 10% may sometimes understate the extent of individual language-selective regions within group-constrained functional parcels.

### Language-selective areas are functionally consistent within individuals

We next assessed the extent to which brain areas, defined by level of language selectivity, are consistently localized (i.e., test-retest reliability as a function of language selectivity). We performed multi-voxel pattern analysis (MVPA) by computing the fMRI response consistency (Pearson’s correlation) of all voxels by decile between the two runs (Haxby et al., 2001). Across anatomical regions, activation pattern consistency as a function of language selectivity always appeared convex, with the most strongly correlated response patterns found among voxels in the most (or least) language-selective deciles, but with inconsistent (uncorrelated) response profiles outside of the areas of peak selectivity (**Fig. 2E–4E**). Using a linear mixed-effects model with polynomial contrasts, we found that the relationship between response pattern consistency and fROI decile was significantly quadratic across all regions examined (*β* > 0, *p* < 0.001; **Supplementary Table 2**), confirming stronger test-retest reliability of voxelwise activation patterns with increasing language selectivity, with the strongest reliability among the most language-selective voxels.

In all lobes, response patterns in the most language-selective deciles were much more consistent than in the intermediate deciles. In the frontal and parietal lobes, increased consistency was also seen for the least language-selective deciles, which may reflect consistency in the location and response of the multiple demand network, which is strongly responsive to the reverse language-localizer contrast (degraded > intact speech; Mineroff et al., 2018). In the least language-selective voxels of the temporal lobe, we did not see such response consistency, paralleling the lack of response to the reverse contrast in the temporal vs. other lobes (**Fig. 2B–4B**).

### Language-selective areas are spatially consistent within, but diverse across, individuals

In addition to the consistency of the fMRI response pattern, we analyzed the spatial consistency of fROI *location* by quantifying the between-run overlap using the Jaccard index (intersection / union of all voxels in the fROIs) by decile. Between-run spatial consistency of voxels of varying degrees of language selectivity was a similarly U-shaped curve across the six lobes (**Fig. 2F–4F**), with significantly greater test-retest spatial consistency among 90%ile language-selective decile voxels relative to the 80%ile (*p* < 0.001; **Supplementary Table 3**). Further, the most language-selective deciles tended to comprise more contiguous clusters of voxels (**Supplementary Fig. 8**), indicating that the more spatially overlapping decile fROIs were also more area-like in their contiguity as opposed to the intermediately language-selective fROIs which were more distributed (less contiguous) and variable in spatial location.

We also quantified the degree of between-subject overlap of language areas in a common (MNI) space, comparing the location of the language-selectivity deciles across all pairs of individuals. High interindividual variability in language fROI location would underscore the importance of precision neuroimaging approaches and suggest the ontogeny of these regions is related to factors other than just their position with respect to cortical folding (Petersen et al., 2024). We found that the spatial consistency across individuals tended to be greater for the 90%ile fROI relative to the 80%ile, however this effect was only significant in the left and right temporal lobes (*p* < 0.01; **Supplementary Table 4**).

Despite the relatively greater between-individual overlap among highly language-selective areas, the likelihood of any two subjects’ language areas being in the same place in a common space was still quite low, especially for the frontal lobe. While the most- and least-selective regions were more likely to overlap across subjects (vs. intermediate “agnostic” areas), the degree of their overlap is low, such that for the most language-selective (90%ile) areas, the vast majority of voxels have an overlap value lower than 60%. This infrequent overlap in language-selective decile fROIs across individuals is shown on the *fsaverage* template (**Fig. 2H– 4H**). The probability of overlap for the top decile language areas was significantly lower across subjects than it was within a subject (*p* < 0.01; **Supplementary Table 5**). These results underscore the high degree of spatial variability between individuals in the precise location of language-selective cortex relative to the standard space (Lipkin et al., 2022). Furthermore, the overlap maps for the “language-agnostic” areas (e.g., 60%ile) illustrate just how rarely the locations of regions of intermediate language selectivity overlap across individuals (**Supplementary Fig. 9**). This finding supports the conclusion that, outside of the most- and least-selective regions, response to language is minimal and spatially inconsistent, both within and between individuals.

## Discussion

### Cortical selectivity for language processing is highly focal

Using fMRI to identify high-level receptive language areas reveals that, in individual brains, the nodes of this network are highly localized and distinct in the degree of their selectivity for language relative to surrounding tissue. This pattern of consistent, focally selective areas provides evidence against alternative, non-modular organizational schemes in which selectivity for language either lacks clear functional boundaries (**Fig. 1A**) or is widely distributed across the cortex (**Fig. 1B**). Our results align with prior work from direct electrical stimulation, in which language-related effects were limited to focal sites (Ojemann et al., 1989; Penfield & Jasper, 1954), as well as from resting state fMRI studies, in which clustering on the basis of functional connectivity patterns also delineated discrete language areas (Braga et al., 2020; Kong et al., 2019; Du et al., 2024; Salvo et al., 2025).

Along with the distinctly increased fMRI responses to language, these focal language areas also had greater spatial contiguity (**Supplementary Fig. 8**) and better within-individual test-retest response consistency (**Fig. 2D-4D, Supplementary Figs. 1D-6D**) relative to adjacent areas of the same size. The functional and spatial distinctiveness of these language-selective regions is consistent with the key functional and structural criteria for “cortical areas” (Eickhoff et al., 2018; Kaas, 1987; O’Leary et al., 2007; Petersen et al., 2024; Rakic et al., 1988) and consistent with a modular organization of language (vs. other cognitive faculties) in the human brain. These findings also align with prior functional connectivity work, in which language network regions identified with a localizer scan were shown to form a consistently co-active core network during language processing, distinct from transiently involved peripheral regions (Chai et al., 2016).

### The most language-selective areas are also the most language-specific

The most language-selective areas were also most suppressed by the spatial working memory task, consistent with prior work suggesting diametric language and “multiple demand” networks (Blank & Fedorenko, 2017; Fedorenko et al., 2012). However, these findings expand upon existing evidence of language modularity by showing that selectivity for spatial WM tended to mirror the selectivity for language, across the language selectivity spectrum, such that the least language-selective regions were distinctly most responsive to the spatial WM task. Moreover, neural response to the spatial working memory task also showed more gradient responses, with voxels in the intermediate deciles exhibiting linear changes in their response (**Supplementary Fig. 7**), suggesting that highly focal areas may be unique to a subset of high-level cognitive faculties, including language, instead of a shared organizational principle for all high-level cognitive faculties. Importantly, in examining the entire cortex, we did not find any evidence that any part of the brain is simultaneously responsive to both language processing and spatial WM, even at low response magnitudes. This finding is in line with other evidence that the neural systems that support linguistic processing vs. those that support non-language cognitive tasks like spatial working memory are highly dissociable (Brunswick et al., 2010; Julian et al., 2019; Kemmerer & Tranel, 2010).

In contrast, the relationship between language selectivity and response to the verbal WM task was more variable across the brain. Sensitivity to verbal WM load was found in some highly language-selective areas but not others, suggesting that any cognitive resources shared between verbal WM and language processing (Acheson & MacDonald, 2009; Ishkhanyan et al., 2019; Schwering & MacDonald, 2020) may depend on a subset of the language network nodes, particularly in parts of STG and IFG (Amici et al., 2007; Fedorenko et al., 2011). Further understanding the nature of these shared neural resources could inform the study of language development and disorders, as verbal WM and language deficits often co-occur in children with developmental communication disorders (Archibald & Harder Griebeling, 2016; Alt, 2011; Larson et al., 2022).

### Regional similarities and variability

Across the frontal, temporal, and parietal lobes in both hemispheres, there was a consistent pattern of uniquely heightened language selectivity within only the top 5-20% of voxels. The shape of these language response profiles showed some lobe-specific variation, but the overarching trend revealed that the 90%ile language fROI was (i) distinctly responsive to language, (ii) distinctly non-responsive to spatial WM, (iii) most reliably localized within individuals, and (iv) most consistent in its voxelwise response pattern across runs.

The temporal lobe had the largest proportion of language-selective tissue, whereas the parietal lobe had the smallest. The focally language-selective area of the parietal lobe was located close to the temporo-parietal boundary (**Fig. 4A,H**) and contiguous with posterior temporal language areas (**Extended Data Fig. 1A**), highlighting how macroanatomical boundaries, especially those that may be less clearly defined like the temporo-parietal boundary (Bokde et al., 2005), may not correspond well with boundaries of the functionally-selective areas (Doherty et al., 1999; Frost & Goebel, 2012). Another example of the macroanatomical and functional misalignment is evident in IFG, where the most language-selective area was found to span contiguous portions of pars opercularis, pars triangularis, and posterior IFS, which atlases commonly parcellate into distinct regions (**Supplementary Fig. 1A,H**). This mismatch may help explain discrepancies in defining Broca’s area (Fedorenko & Blank, 2020; Tremblay & Dick, 2016) and the high degree of discord across studies that have attempted to functionally subdivide the IFG based on group-average data (Regev et al. 2024; Siegelman et al., 2019).

Language selectivity in the temporal lobe, including STG and MTG, covered more area but was more distinctly circumscribed and more consistently localized both within and between individuals relative to the frontal and parietal language areas. This may be partially attributable to the auditory modality of the language localizer employed in this study, to speech as the native language modality in our subjects, or to developmental primacy of the temporal lobe for language. Temporal regions are closer to primary auditory areas, which may constrain the development of their functional specialization according to the tethering hypothesis (Buckner & Krienen, 2013; Wang et al., 2023). This may also reflect a more “primary” role of temporal language areas in the ontogeny of the language network (Skeide & Friederici, 2016), in line with a case study that found frontal language selectivity is abnormally non-focal in the absence of temporal language areas (Tuckute et al., 2022).

### Hemispheric similarities and differences

Our characterization of the distinct focality of language areas was similar for canonical left hemisphere language areas and their right hemisphere homologues, including in terms of functional selectivity, between-run consistency in location and activity, and spatial contiguity. In line with previous reports of language being left-lateralized in the majority of individuals (Lipkin et al., 2022; Loring et al., 1990; Martin et al., 2022), the language fMRI response magnitude and between-run consistency measures were typically greater in the left hemisphere. The similar pattern of findings bilaterally accords with prior work that found language to be organized similarly in both hemispheres (Martin et al., 2022; Quillen et al., 2021) and may reflect the right hemisphere’s early potential to support language that decreases throughout development (Berl et al., 2014; Holland et al., 2007; Lenneberg 1967, 1969; Olulade et al., 2020; Peña et al., 2003; Tuckute et al., 2022). Future work is needed to decipher whether this similarity in functional organization reflects similarity in neurocomputational contributions to language processing.

### Within-individual consistency and between-individual variability in localization

In all anatomical regions examined, there was a pattern of increased variability, both within and between individuals, in the location and activity patterns of areas with intermediate levels of language selectivity relative to the more consistently located minimally and maximally language-selective fROIs. This pattern of disorganized, inconsistent, minimal response to language is incompatible with models of distributed or gradient language selectivity outside of core functional language areas. Given the ability to resect vast sections of the frontal and temporal lobe without impacting language ability (Rasmussen & Milner, 1975), it may be that these intermediate regions are wholly inessential to high-level language processing. Alternatively, they may be only transiently involved in language processing (Chai et al., 2016; Fedorenko & Thompson-Schill, 2014) in ways that are peripheral to the core computations involved in combinatorial meaning, such as accessing specific concepts in semantic or episodic memory, which are known to be distributed across the cortex (Farah et al., 2004; Huth et al., 2016). Further work is needed to determine what features, if any, elicit activation in non-core language areas during language processing and what the timecourse and functional consequences of these activations are (Regev et al., 2024).

Although the location of the most language-selective areas in a standard space is variable across individuals, both in the current data and in prior reports (Mahowald & Fedorenko, 2016; Ojemann et al., 1989), the organization of less-selective regions appears to be essentially random both within and across individuals. This further reinforces the conclusion that there is no systematic functional language gradient outside of focal nodes. Moreover, that there is at least some consistency in the localization of maximally language-selective areas across subjects suggests there may be yet-undiscovered principles guiding the organization of the human brain’s language areas beyond merely their location with respect to the cortical curvature. For instance, functional cortical areas can often be localized with respect to unique patterns of thalamocortical connectivity, cytoarchitecture, and myeloarchitecture (Cadwell et al., 2019; Fenlon & Suárez, 2013; O’Leary et al., 2007; Passingham et al., 2002; Petersen et al., 2024; Rakic, 1988). Better characterization of these microstructural features *in vivo* may allow us, in future work, to predict the location of focal language areas in individual brains with greater precision than relying on coarse gyrus- or atlas-level descriptions.

### Implications for the functional localization approach

Defining single-subject fROIs by applying a fixed, *a priori* threshold (e.g., the top 10% of voxels) to statistical parametric maps (Fedorenko et al., 2010; Julian et al., 2012) has become a popular approach to investigating not only the language network (e.g., Fan et al., 2024; Lee et al., 2024; Martin et al., 2022; Ozernov-Palchik et al., 2026), but also regions implicated in other cognitive domains including high-level vision (Molloy et al., 2024) and social interaction (Isik et al., 2017). The stated motivation for a fixed threshold has been to have a consistent fROI volume across individuals. However, this assumption requires drawing fROIs in a standard space like the MNI or fsaverage templates, which distorts functional boundaries through interpolation relative to building fROIs in subjects’ native space. Furthermore, we have shown that the relative extent of language-selective cortex may differ across the brain’s lobes and hemispheres, with a much larger proportion of language-selective voxels in the temporal than frontal lobe. Importantly, here we also show that a single *a priori* threshold can obscure differences in the size of functional cortical areas across subjects. We resolve these discrepancies by introducing the novel approach of localizing single-subject fROIs on the basis of sharp transitions in the magnitude of individual activation profiles (i.e. the point of maximum curvature in the ranked response distribution). This individualized, data-driven approach better reflects known variability in the size of functionally specialized areas across individuals (Caspers et al., 2006; Dougherty et al., 2013; Dworetsky et al., 2024; Gao et al., 2022; Gordon et al., 2017; O’Leary et al., 2007). Such variability may be relevant to behavioral outcomes (Leingärtner et al., 2007), as well as disorders in which the cortical arealization may be altered (e.g., polymicrogyria; Lenge et al., 2019). More individualized approaches in neuroscience may help improve generalizability of findings and translational relevance (Nebe et al., 2023; Seghier & Price, 2018). This approach could also help validate whether other cognitive faculties are suitable candidates for the functional localization approach. Just because core receptive language in the brain is associated with several highly focal nodes does not mean that all cognitive operations have this kind of organization (Westlin et al., 2023). Functional activation maps that lack such sharp transitions in their activation profiles may reveal that other brain processes are distributed or gradient and therefore not appropriately characterized by using circumscribed single-subject fROIs.

Although fROIs defined with the individualized, data-driven approach did yield different language-fROI sizes across individuals, these fROI thresholds were often close to the commonly employed “top 10%” threshold, validating this methodological choice in prior work. Despite the similarity in size of both sets of fROIs, knee-based fROIs may have starker functional boundaries with surrounding tissue compared to the top 10% fROIs which tended to be over-inclusive and thus appear less specialized for language (**Supplementary Figs. 10-11**). Using this more personalized localization approach in future fMRI studies involving language in different task contexts could further refine our understanding of language fROIs as cortical areas with clear boundaries (Eickhoff et al., 2018; Petersen et al., 2024).

### Limitations and future work

Functional MRI is an indirect measure and of neural activity, so additional work combining fMRI with more direct recording (e.g., electrocorticography; “ECoG”) or stimulation (e.g., direct electrical stimulation, transcranial magnetic stimulation) of neural activity itself will be important to clarify just how sharp the functional boundaries of language areas are. For instance, ECoG studies often report language-selective electrodes immediately next to non-selective ones (Fedorenko et al., 2016; Woolnough et al., 2023), but correspondence with fMRI-localized language area boundaries is understudied. Additionally, our work was limited to healthy young adult brains, leaving the ontogeny of these circumscribed language areas an important area for future work. Neural development of language areas has been hypothesized to mirror the development of behavioral top-down, higher-level language processing abilities (Skeide & Friederici, 2016), with language-selective areas, particularly in the frontal lobe, emerging only later in development. Further, lateralization of the fMRI-localized language network changes across early childhood (Ozernov-Palchik et al., 2026), which may reflect increasing modularization of language-selective regions, along with the development of these stark functional boundaries.

Additionally, this work primarily focused on identifying the *functional* boundaries of language-selective areas, and future work is needed to clarify whether there exist any corresponding *structural* boundaries that characterize these regions (e.g., in the cytoarchitecture or in patterns of thalamocortical projections; Eickhoff et al., 2018; Petersen et al., 2024). A case study of single individual born without a left temporal lobe suggests that connectivity between language-related structures may be key to typical arealization of language-selective areas (Tuckute et al., 2022). Multimodal MRI studies investigating the correspondence between structural and functional features (e.g., Saygin et al., 2016) could provide further evidence that there are structurally distinct cortical language areas.

Our findings also relate to the phylogeny of language, as ongoing research in the evolution of language examines the extent to which the functional and structural organization of communication-related processing differs in non-human primates (Changeux et al., 2021; Friederici & Becker, 2025; Skeide & Friederici, 2015). Many language-associated macroanatomical features are relatively evolutionarily conserved across non-human primates (e.g., structural left-lateralization of planum temporale; Friederici, 2026). However, in light of the interindividual variability and focality of language areas in humans, the study of the evolutionary origins of language areas may benefit from more precise characterization of language-associated structures (which often deviate from common macroanatomical landmarks).

### Conclusions

Using precision fMRI, we found that, in individual brains, cortical areas for high-level receptive language processing are highly focal and sharply circumscribed, rather than broadly or gradiently distributed across the cortex. These language-selective areas are uniquely and strongly responsive to language and, across repeated measurements within an individual, highly consistent in their functional response pattern and spatial location. Characterized by a sharp transition in language response magnitude, these areas are also variable in size and location across individuals. Taken together, the functional properties of receptive language areas are consistent with the functional criteria for “cortical areas” in systems neuroscience (Cadwell et al., 2019; Eickhoff et al., 2018; O’Leary et al., 2007; Passingham et al., 2002; Petersen et al., 2024; Rakic, 1988). This raises the possibility that, like other cortical areas, human language areas may also be characterized by unique structural features that distinguish them from adjacent cortex. Future work to identify these features will help reveal the ontogenetic and phylogenetic origins of this uniquely human ability.

## Materials and Methods

### Participants

Twenty-eight participants (16 female, 12 male; age: mean = 22.9 years, range = 19-32 years) recruited from the Boston area completed structural and functional imaging for this study. The Institutional Review Board of the Boston University Charles River Campus and the Massachusetts Institute of Technology Committee on the Use of Human Subjects as Experimental Subjects approved this study. Participants were fluent American English speakers with no history of difficulties with speech, hearing, reading, language, or cognitive and motor development.

### In-Scanner Tasks

Participants completed three in-scanner tasks: a *language localizer* (Scott et al., 2017), a *verbal working memory task* (Fedorenko et al, 2011), and a *spatial working memory task* (Fedorenko et al, 2011; Lee et al., 2024). During the language localizer, participants passively listened to either intact or unintelligible, degraded speech. Stimuli consisted of audio clips from engaging media including speeches, interviews, and podcasts. Two runs of 5min 58sec were performed with each comprising 8 intact blocks, 8 degraded blocks, and 5 rest blocks where participants fixated on a “+” and no other stimuli were presented. Degraded stimuli were produced by low-pass filtering the natural recordings and adding white noise modulated with the original amplitude envelope, rendering these stimuli completely unintelligible (Scott et al., 2017). Sound clips used in the intact and degraded blocks were distinct such that a participant did not hear the same clip in both conditions. Intact speech trials were contrasted with degraded speech trials (intact > degraded) to determine voxelwise selectivity for language within individual subjects.

The verbal working memory (VWM) task was derived from classic digit span tasks. Participants heard two sequences of digits and pressed a button to indicate if the sequences matched. Sequences were either 3 digits long (low verbal working memory load) or 6 digits long (high load). Digit recordings in the first sequence were recorded by a female speaker and those in the second by a male speaker (to emphasize the abstract, verbal content rather than sensory memory for the stimulus). Mismatch trials involved transposition of a single adjacent pair of digits (e.g., “5-8-3-1-2-6”; “5-8-3-2-1-6”). In each run, there were 5 blocks of each of the two conditions. Activation in the 6-item (high load) condition was contrasted against the 3-item (low load) condition to identify fMRI activity related to sensitivity to verbal working memory load.

The spatial working memory (SWM) task used was derived from the classic Corsi block-tapping test (Corsi, 1972). A 3 x 3 grid of gray circles was presented on the display, and then 3 (low spatial working memory load) or 6 (high load) positions were illuminated one-by-one in red in a random, non-repeating sequence. Then, the same number of positions was illuminated in blue. Participants were instructed to indicate whether the two sequences of positions were identical via button press. Mismatch trials involved transposition of a single adjacent pair of locations. As with the VWM contrast, the 6-item (high load) condition was contrasted against the 3-item (low load) condition to identify voxelwise sensitivity to spatial working memory load. A more detailed account of the fMRI task paradigms is available in Scott (2020).

### Data Acquisition

Structural and functional MRI scans were collected at the Athinoula A. Martinos Imaging Center at MIT with a Siemens Trio 3T scanner and 32-channel head coil. The structural scans were acquired with a T1-weighted (T1w) magnetization prepared rapid gradient echo (MPRAGE) sequence [TR = 2530 ms, TE = (1.64, 3.50, 5.36, 7.22 ms), TI = 1400 ms, flip angle = 7.0°, voxel size = 1.0 mm isotropic, FOV = 256 x 256 mm, 176 sagittal slices] and a T2-weighted sequence [TR = 3200 ms, TE 454 ms, voxel size = 1.0 mm isotropic, FOV = 256 x 256, 176 sagittal slices]. For the functional volumes, a simultaneous multislice T2*-weighted gradient EPI sequence with continuous sampling was used [TR = 750 ms, TE = 30 ms, flip angle = 90°, voxel size = 3.0 mm isotropic, 10% slice gap, FOV = 72 x 72, 45 slices, 5 simultaneous slices].

### Data Analysis

#### Preprocessing

Anatomical T1w volumes were processed with FreeSurfer’s (v5.3.0) cortical reconstruction workflow (Dale et al., 1999), using T2w volumes to improve estimation of the pial surface. Functional data were processed with Nipype v0.13 workflows (Gorgolewski et al., 2011) in the Lyman platform v1.0 (Waskom, 2019). Preprocessing consisted of within-run motion correction and detection of outlier volumes. Within-run correction was performed with rigid-body realignment to the mean EPI image with FMRIB Software Library’s (FSL; Jenkinson et al., 2012) *MCFLIRT*. Volumes were considered outliers if the framewise motion exceeded 1 mm or the global signal intensity was more than 3 standard deviations from the run mean. Outlier volumes and the 6 motion parameters produced by *MCFLIRT* were included as nuisance regressors in the general linear model (GLM; Siegel et al., 2014). The functional data used for the following analyses were not spatially smoothed (because we wanted to examine how focal or graded the task-fMRI responses were without any added smoothness), but similar results were obtained using smoothed data (**Fig. S12).**

#### Single-Subject Modeling

Two task regressors for each contrast of interest were included. These task regressors were generated by convolving a vector of event onsets with the event durations. Then, this stimulus time series was convolved with the canonical hemodynamic response function to generate the hypothesized BOLD response. Within-subject estimation of the GLM and the contrasts of interest were completed for each run with volumes in subjects’ native EPI space. Volumes with z-statistic and parameter estimate maps from the GLM were then transformed to the coordinate space of subject’s T1w volume using FreeSurfer’s *tkregister* and *mri_vol2vol*.

Functional ROI-based analyses were performed for 9 regions in each hemisphere, the frontal, parietal, and temporal lobes, as well as two smaller regions per lobe from the Desikan-Killiany parcellations (e.g., inferior frontal gyrus, middle temporal gyrus). We also performed supplementary analyses with Group-Constrained Subject-Specific parcels (Fedorenko et al., 2010; Julian et al., 2012) for comparability with the many papers that have defined fROIs with similar fMRI data-driven parcels. Binary anatomical volumes were isolated from each individual’s anatomical parcellation for frontal, temporal, and parietal regions, and the following analyses were all performed in the individual’s anatomical T1w coordinate space.

The *z*-statistic map indicating activation significance for the language localizer contrast (intact > degraded) was used to define percentile-based functional ROIs (fROIs) within each anatomical region. First, the z-statistic map was masked with the target anatomical region (e.g., left frontal lobe, left IFG). Deciles of the values in the masked z-statistic map were calculated. The masked z-statistic map was then thresholded and binarized to create a map of voxels for which the z-statistic was within each decile range (0-10th percentile, 10-20th percentile, …, 90-100th percentile). This procedure resulted in 10 binary fROI volumes per anatomical ROI for each individual.

The average response within the language localizer-defined fROIs was calculated for the language contrast (intact > degraded), SWM contrast (high > low), and VWM contrast (high > low). For each task, the contrast parameter estimate volume was masked with the language fROI volume, and the nonzero average of the masked parameter estimate was computed. The data used to define the fROI and the data used to calculate the average response were from different runs. For the VWM task, one subject fell asleep during volume acquisition and was excluded from all analyses involving the VWM contrast.

#### Activation-decile fROIs

For each anatomical ROI, the average language localizer response within the language activation-decile fROIs was modeled using a linear mixed-effects model with the R package lmerTest (v3.1-3). The dependent measure was the average language localizer response for each run and each individual. The decile (0-10^th^, 20-10^th^, …, 90-100^th^ percentile) was included as a fixed factor with 10 levels with polynomial contrast coding. The participant was the random effect. The function *lmer* was used with Restricted maximum likelihood (REML) estimation for model fitting.

#### Data-driven within-subject language-selectivity thresholding

For each individual, the average fMRI response per decile fROI was also computed again at a more granular level for the language localizer contrast. The voxels with activation significance in the top 25% were assigned to single percentile interval fROIs: 75^th^-76^th^ %ile, 76^th^-77^th^ %ile, …, 99^th^-100^th^ %ile. The average fMRI response (averaged across the two runs) was plotted against the percentile interval index, and the knee of each curve, a local maximum in terms of curvature, was identified. The *KneeLocator* function from the kneed python package (v0.8.5; Satopa et al., 2011) was used to fit splines to the data with polynomial interpolation, and the knee was located under the assumption that the curve was convex and increasing. This approach failed to find a knee for 14 of 168 total curves. For 7 of these, the knee was found by first fitting a different model to the data, a linear combination of exponential functions, using SciPy’s (v1.11.1) *curve_fit*, then using *KneeLocator*. For the other 7 curves, there were not clear knees because the curves were not convex, and these individuals were excluded from the boxplots of knee locations (**Figs. 2D–4D, Supplementary Figs. 1D–6D**).

#### Between-run multi-voxel pattern analysis (MVPA)

Multivoxel pattern analysis (MVPA) of the response profile within each fROI were conducted to quantify the test-retest reliability for the language localizer contrast and the similarity in response profile across tasks. The Pearson’s correlation coefficient for the fROI-masked response volumes of the language localizer contrast across the two runs was calculated. This was performed twice, swapping out the fROI used (defined from run 1 data vs. run 2 data), and these subject-specific correlation coefficients were averaged. Then, for each decile, the average correlation coefficient across all subjects was computed. The LME modeling procedure used for language localizer response across fROI deciles was used again here with correlation coefficient as the response variable.

#### Spatial contiguity of language-selective fROIs

The final intra-individual spatial fROI analysis concerned the degree of connectedness within each decile fROI. The *connComp3D* function from the R package neuroim (v0.0.6; Buchsbaum, 2016) was used to identify connected groups of nonzero voxels in the binary fROI masks. Connected voxels were identified by considering the 26-voxel neighborhood of each nonzero voxel and assigning any nonzero neighbors to the same cluster as the current voxel. The weighted mean cluster size for each decile was calculated by multiplying the proportion of the total fROI voxels in each cluster by the cluster size.

#### Anatomical consistency of language-selective regions

In addition to the within-subject language localizer response profile similarity across runs, the spatial similarity between each pair of run-specific decile fROIs was calculated. For each subject, the Jaccard index for all the decile- and region-specific fROIs was calculated using FSL’s *fslmaths* to multiply the binary decile fROI volumes from each run and *fslstats* to count the nonzero voxels in the output overlap map. The number of overlapping voxels was then divided by the total number of nonzero voxels across the two fROI volumes to get the Jaccard index. These indices were averaged across all subjects for each decile of each anatomical ROI.

#### Between-subject consistency of language-selective fROIs

Some inter-individual analyses and visualizations required individuals’ data to be in the same coordinate space. The contrast z-statistic volumes were normalized to MNI coordinates by first registering the subject’s native EPI volume to the T1w volume and then warping the T1w volume to MNI space. FreeSurfer’s *BBRegister* (Greve and Fischl, 2009) with *FLIRT* initialization was used to rigidly register the subject’s native mean EPI volume and the FSL MNI152 template (v5.0.7). Next, the subject’s T1w volume was mapped to the MNI template using ANTs’ (v1.9; Avants et al., 2011) *antsIntroduction.sh* with default parameters used to map the subject’s T1w volume to the MNI template. These transformations were then applied to the GLM output volumes using FreeSurfer’s *mrivol2vol* and ANTs’ *WarpImageMultiTransformation*. This was performed for each of three contrasts of interest: language localizer (intact > degraded), SWM (high load > low load), VWM (high load > low load). The following analyses involved only these normalized volumes.

The fROI delineation procedure described above was repeated in MNI coordinates. First, each subject’s Desikan-Killiany parcellation volume was normalized to MNI coordinates using ANTs’ *WarpImageMultiTransform* and the transformation matrix and deformation field generated during the normalization of functional data. The decile-specific fROIs were again defined for each of the 18 anatomical ROIs using normalized z-statistic volumes.

To investigate the spatial consistency of decile fROIs across individuals, the between-subject fROI Jaccard index was also calculated pairwise for each decile of each anatomical region. The Jaccard index was calculated for all possible pairs of subjects, using fROI volumes from the same run. The by-run indices for each subject-pair were first averaged. Then, the average Jaccard index for each anatomical region decile was computed.

To investigate if the within-subject and between-subject overlap was greater for the 90%ile fROI compared to the 80%ile fROI, and to see if this increased overlap was greater for the within-subject measure, we used a linear mixed effects model. For each decile and each region, the relevant data for each subject included the between-run overlap (run 1 vs. run 2) and the average of the between-subject overlap values for all subject pairs including the subject of interest. Decile (80%ile, 90%ile) and measure (within-subject JI, between-subject JI) were fixed effects, and we included the interaction of these two effects. Subject was included as a random intercept due to the two measures being related (i.e., the within-subject JI and between-subject JI may be correlated. Bonferroni correction was used to correct for the 18 tests run (one per anatomical region).

To visualize fROI locations across deciles and subjects, fROI probability maps for each decile were generated for the 18 anatomical regions of interest. First, fslmaths was used to sum individual fROI volumes from each of two runs per participant and divide the aggregate volume by twice the number of participants. This probability volume was mapped to the fsaverage surface using FreeSurfer’s *mri_vol2surf*. PySurfer (v0.11; Waskom et al., 2020) was used to depict the overlay on the inflated fsaverage surface.

## Acknowledgments

We thank Jessica Tin, Yaminah Carter, and Ja Young Choi for assistance with data collection; and Atsushi Takahashi, Steve Shannon, and Sheeba Arnold at the Athinoula A. Martinos Imaging Center at the McGovern Institute for Brain Research, MIT, for technical support during scanning. Research reported in this article was supported by the National Institutes of Health (NIH) under award numbers R03 DC014045 and R03 HD096098 to TP. RB was supported by T32 DC01301 and F31 DC022801. TS was supported by T90 DA032484 and F32 DC019531.

