## Supplementary Information for "Selectivity for high-level language processing is highly localized in individual brains"

### Extended Data

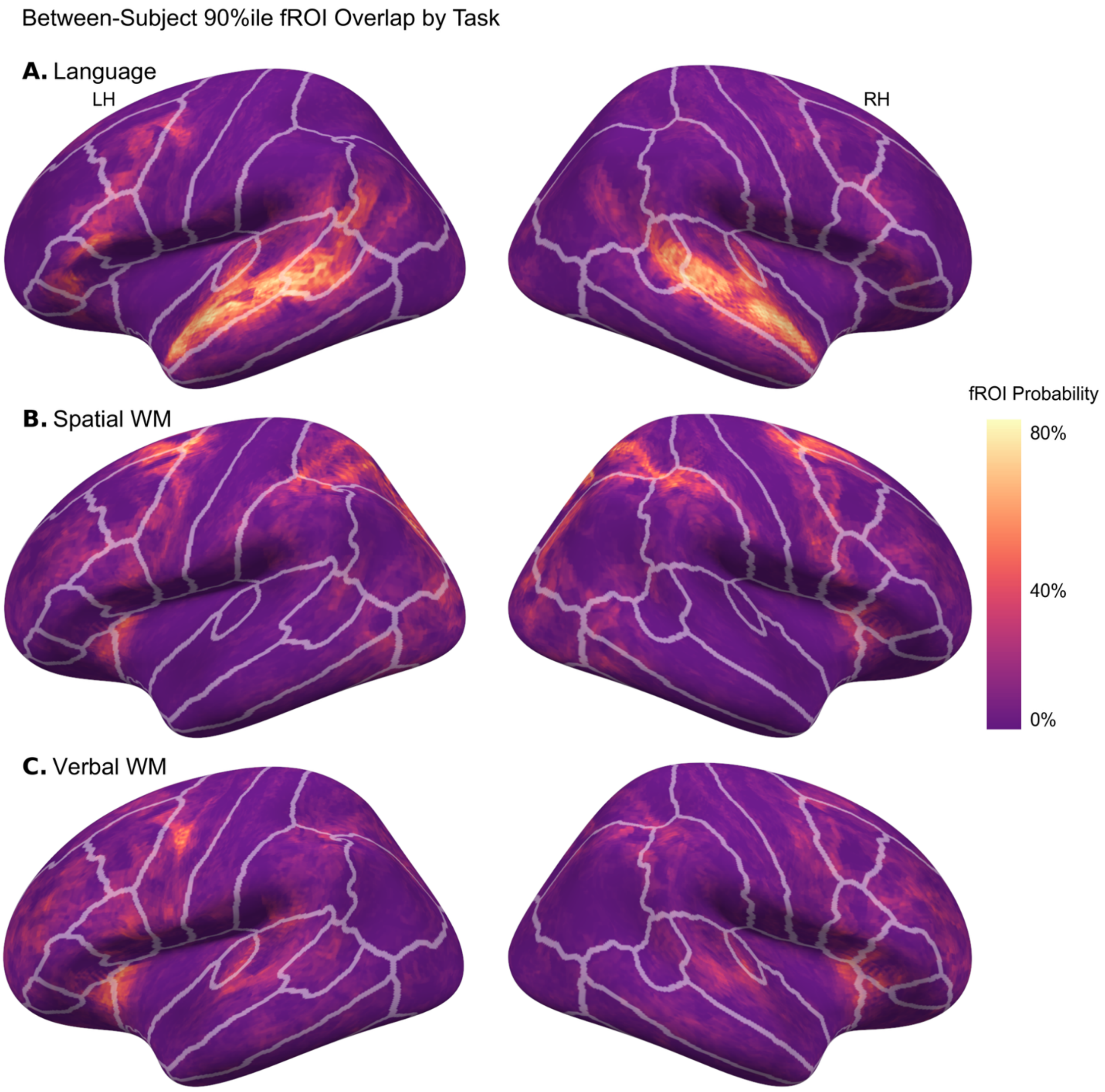


**Extended Data Figure 1**: The probability of each vertex being included in the 90%ile fROIs (based on two run-specific fROIs per subject) for the **(A)** language localizer, **(B)** spatial working memory, and **(C)** and verbal working memory tasks. fROIs were defined within the cortical ribbon for the entire left hemisphere (left column) or right hemisphere (right column). Outlines show Desikan-Killiany ("aparc") atlas regions.

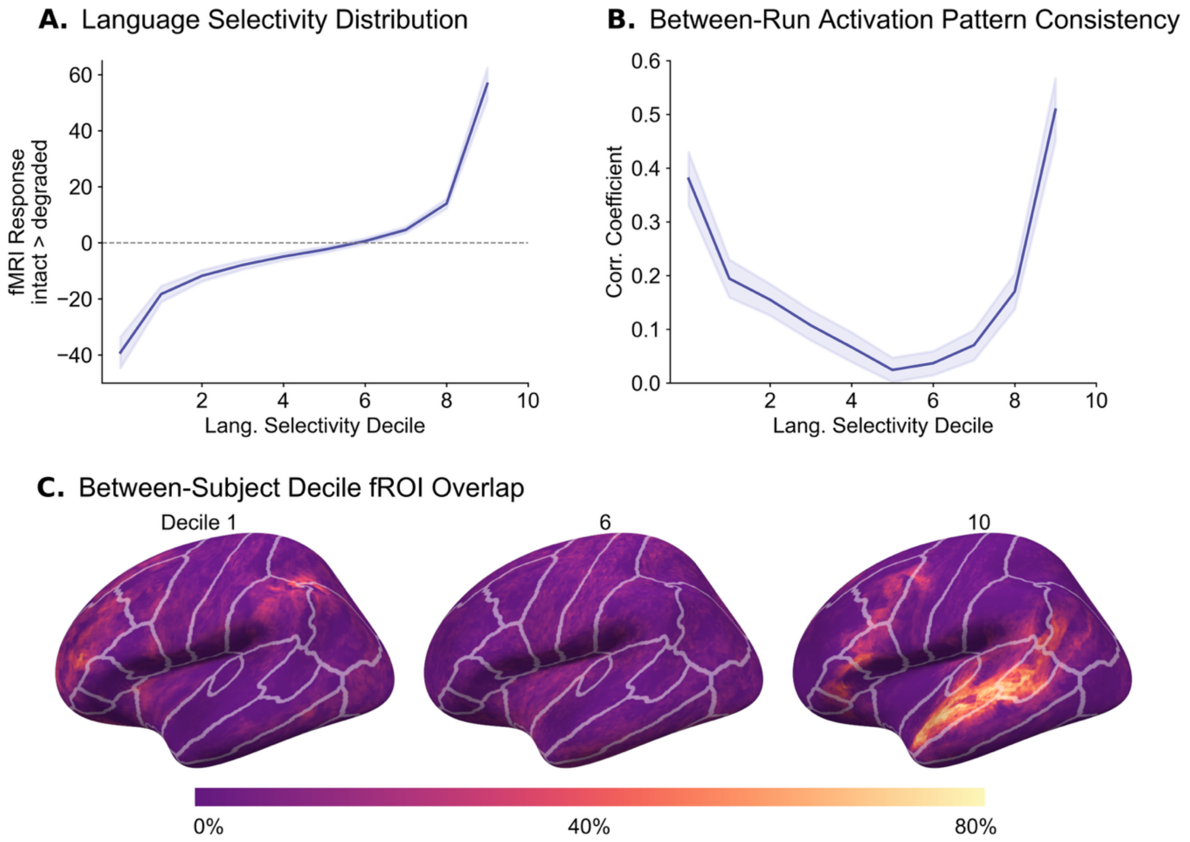


**Extended Data Figure 2**: Across the whole left hemisphere cortex, functional response to language exhibits a highly localized pattern in individual brains. **(A)** The average language response magnitude across subjects as a function of language selectivity (grouped by decile) shows steep decreases in response to language outside of the most selective voxels. **(B)** The average voxelwise correlation across subjects between activation maps from the two runs of the language localizer task as a function of language selectivity (grouped by decile). The most language-selective regions have distinctly heightened test-retest consistency in response to language. **(C)** Probability maps for fROI location, shown for the 1st, 6th, and 10th deciles. The percentage indicates the proportion of subjects with a decile-fROI that includes that vertex. Note that areas of “intermediate” language selectivity are organized essentially randomly across subjects.

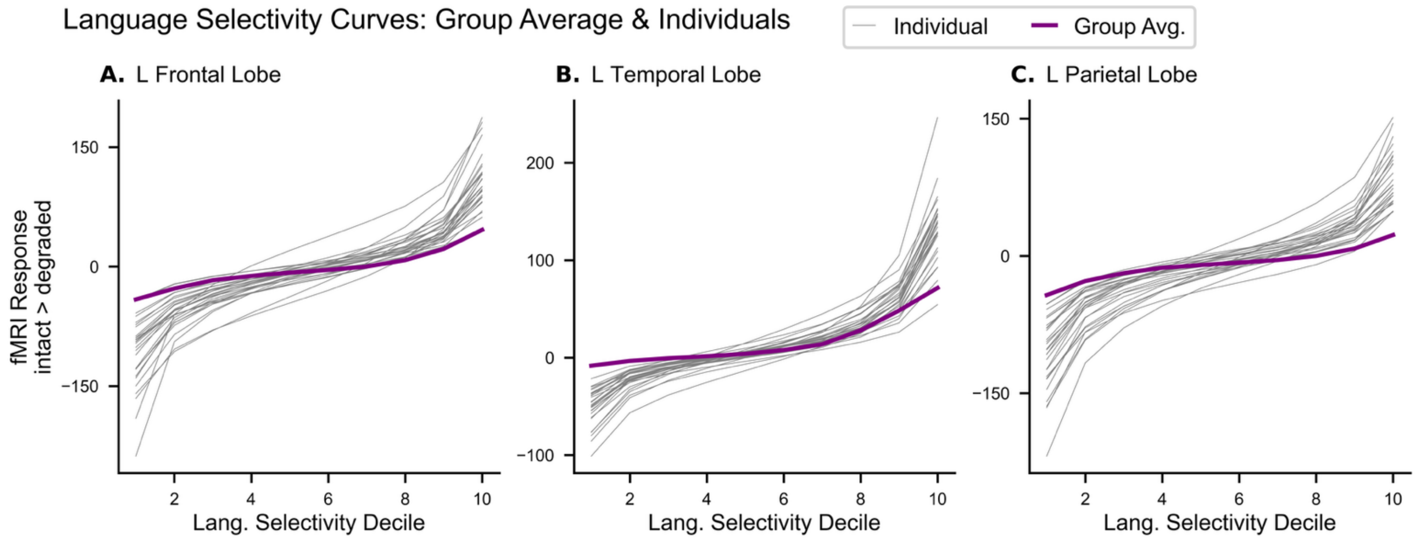


**Extended Data Figure 3**: Individual subjects' language response magnitude as a function of language selectivity (grouped by decile) within the **(A)** left frontal lobe, **(B)** left temporal lobe, and **(C)** left parietal lobe. The results from classical univariate group-average fMRI data in MNI space (purple) and from each individual subject's data in native space (grey) are shown. Linear regression models were fit to both the group-level and individual-level curves, in order to ascertain the extent to which group average data underestimated the focality and overestimated the gradience of language response. $R^2$ values were computed for each model to indicate how well a linear model described the data. Bootstrapping (1,000 samples) was used to get a distribution for the group-level $R^2$. One-sided Mann-Whitney U-tests (one per lobe) indicated that the $R^2$ values were significantly lower for the left frontal and left parietal lobes ($U$ = 9679, $p < 0.005$, $U$ = 7577, $p < 0.005$, respectively) but not for the left temporal lobe ($U$ = 12081, $p > 0.05$). These data show that the focality of language-selectivity is much stronger in individual brains than what would be estimated based on classical univariate group-average approaches.

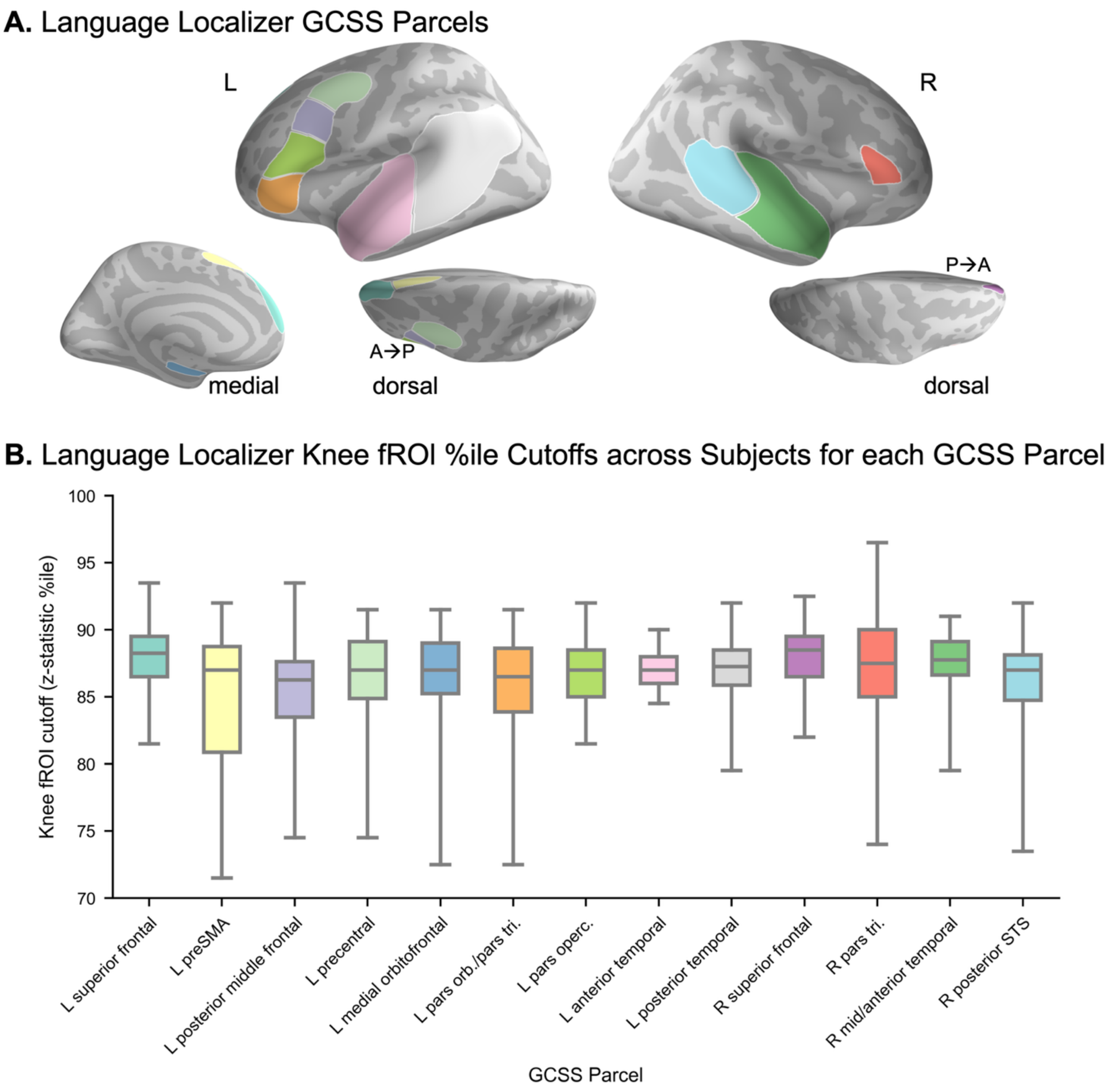


**Extended Data Figure 4:** Individualized fROI approach within functionally (not anatomically) defined search-space parcels. **(A)** Parcels defined using the group-constrained subject-specific (GCSS) approach (Fedorenko et al., 2010; Julian et al., 2012) in which parcels are drawn using a smoothed probability map indicating where, across all subjects, significant activation to the language localizer contrast is likely to be located. **(B)** Boxplots depicting the personalized fROI boundaries (expressed as contrast *z*-statistic %ile) across the 28 individuals for each GCSS parcel. Cutoff percentiles were typically lower (fROI sizes > 10% of parcel) for the GCSS parcels than for the atlas-defined anatomical parcels, as the search space was constrained to areas more likely to have language activation. Moreover, these fROI-forming thresholds tended to be more inclusive than the usual top-10% approach adopted in most prior studies using GCSS, validating the use of that cutoff in prior work in identifying language-selective areas, but also suggesting it may be too conservative if the goal is to capture the full extent of language selective areas.

### Supplementary Information

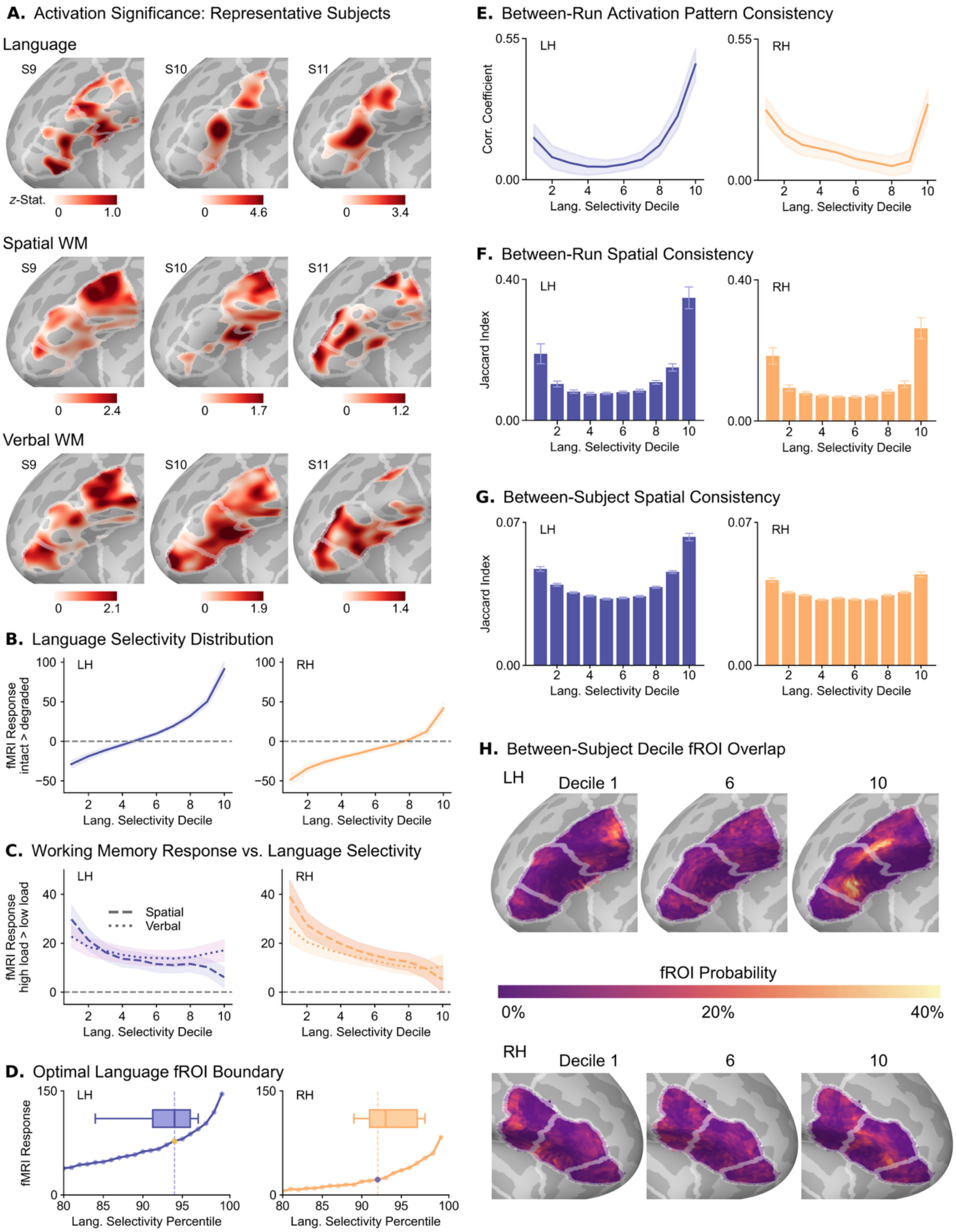


**Supplementary Figure 1**: Inferior frontal gyrus. **(A – C, E – H)** Same descriptions as in Figure 2. **(D)** The knee point for language selectivity at the group average level (average of individuals' fMRI response values for each percentile; dashed lined indicating knee at: LH: 93%ile, RH: 91%ile) and at the individual level (box plot of individuals' knee values; median: LH: 93%ile, RH: 92%ile).

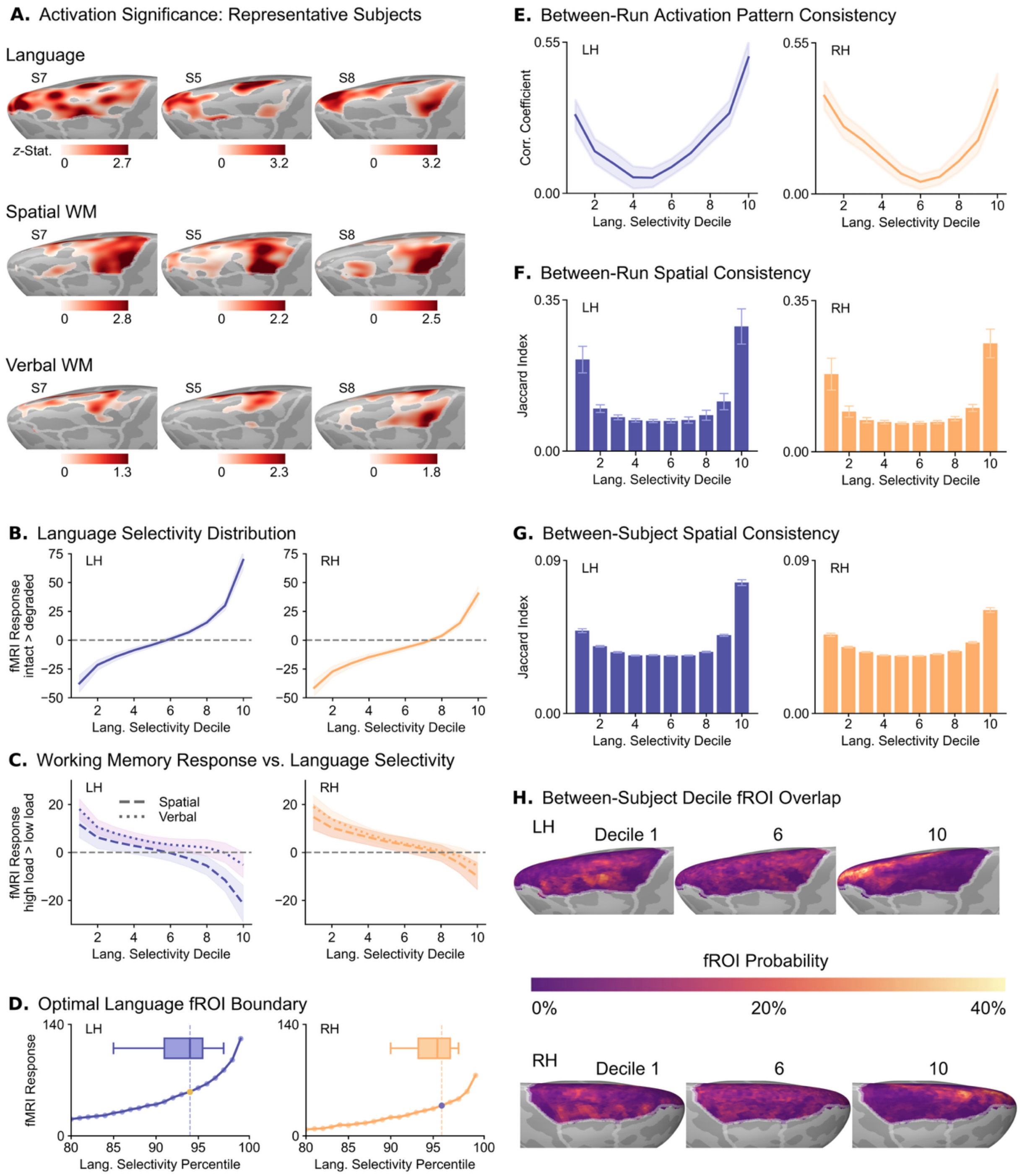


**Supplementary Figure 2**: Superior frontal gyrus. **(A – C, E – H)** Same descriptions as in Figure 2. **(D)** The knee point for language selectivity at the group average level (average of individuals' fMRI response values for each percentile; dashed lined indicating knee at: LH: 93%ile, RH: 95%ile) and at the individual level (box plot of individuals' knee values; median: LH: 93%ile, RH: 94.5%ile).

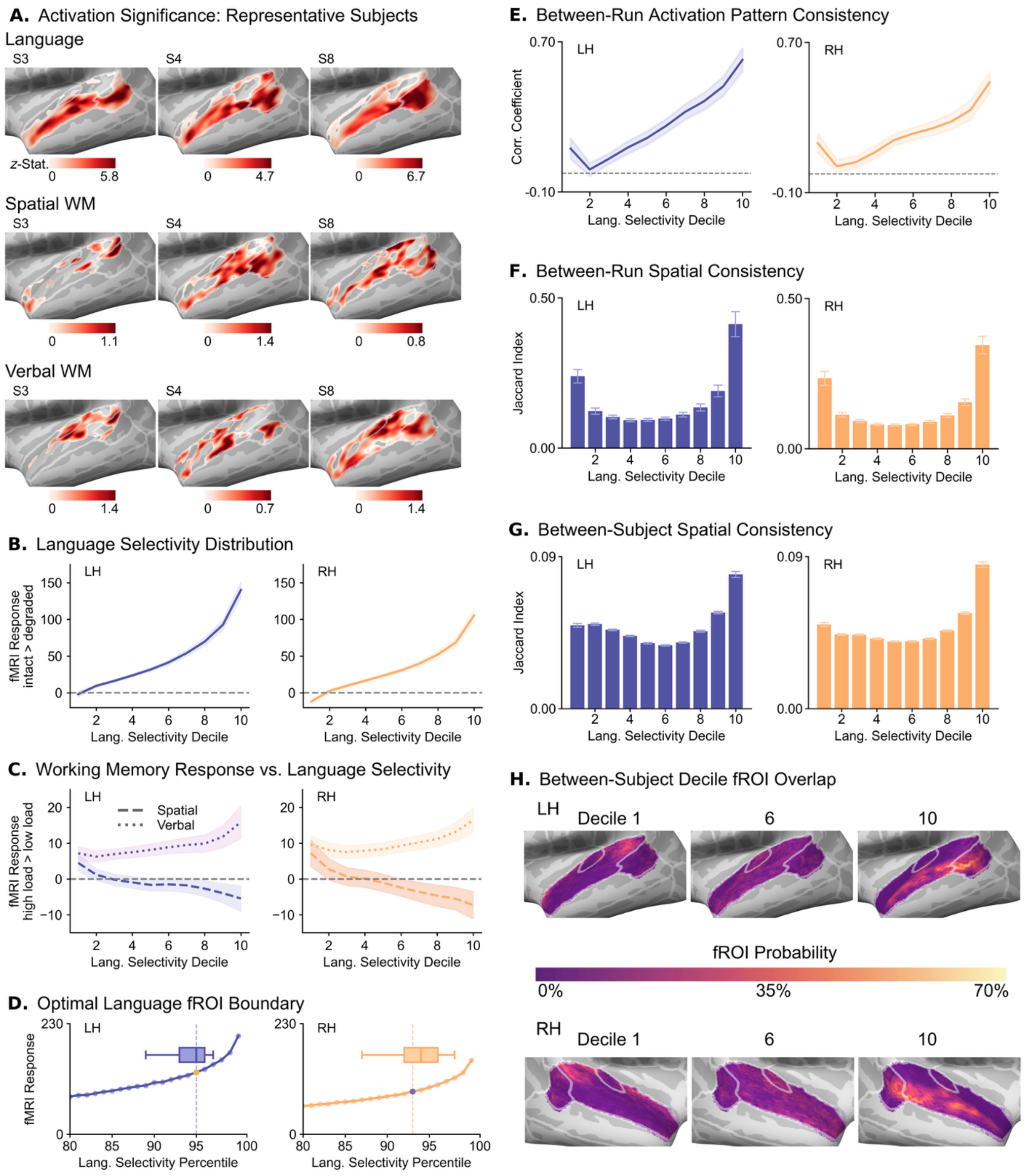


**Supplementary Figure 3**: Superior temporal gyrus. **(A – C, E – H)** Same descriptions as in Figure 2. **(D)** The knee point for language selectivity at the group average level (average of individuals' fMRI response values for each percentile; dashed lined indicating knee at: LH: 94%ile, RH: 92%ile) and at the individual level (box plot of individuals' knee values; median: LH: 94%ile, RH: 93%ile).

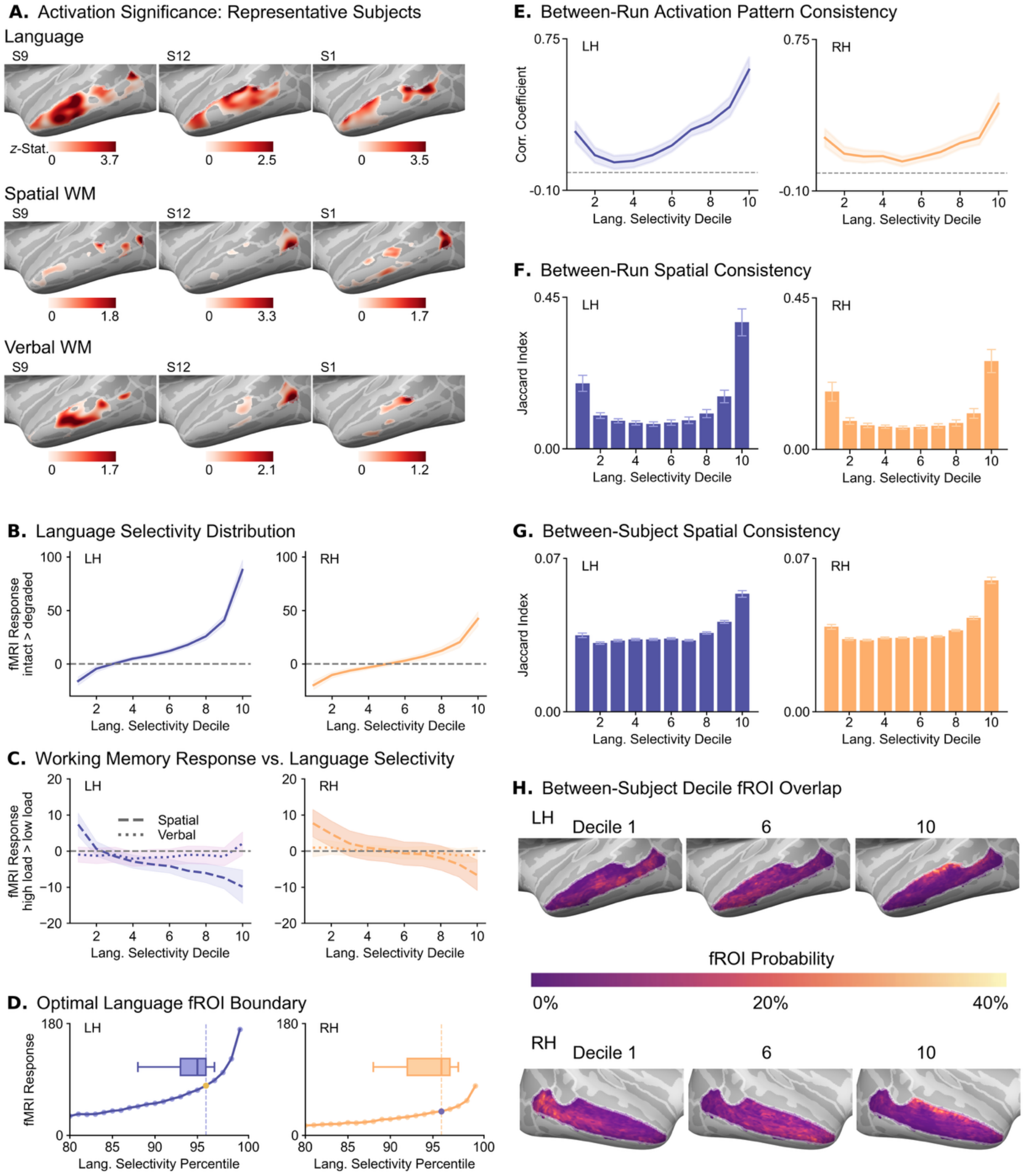


**Supplementary Figure 4**: Middle Temporal Gyrus. **(A – C, E – H)** Same descriptions as in Figure 2. **(D)** The knee point for language selectivity at the group average level (average of individuals' fMRI response values for each percentile; dashed lined indicating knee at: LH: 95%ile, RH: 95%ile) and at the individual level (box plot of individuals' knee values; median: LH: 94%ile, RH: 95%ile).

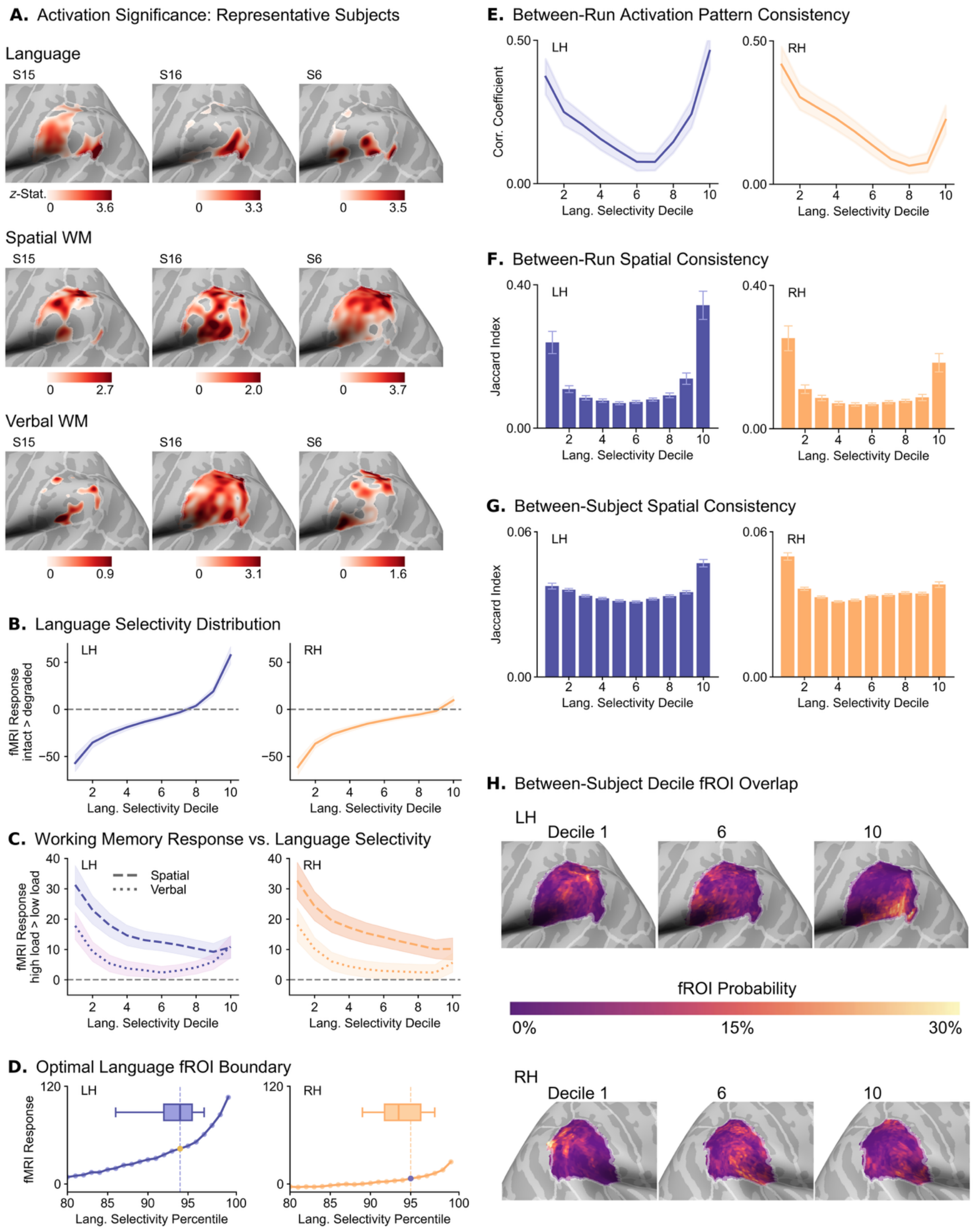


**Supplementary Figure 5**: Supramarginal Gyrus. **(A – C, E – H)** Same descriptions as in Figure 2. **(D)** The knee point for language selectivity at the group average level (average of individuals' fMRI response values for each percentile; dashed lined indicating knee at: LH: 93%ile, RH: 94%ile) and at the individual level (box plot of individuals' knee values; median: LH: 93%ile, RH: 92.5%ile).

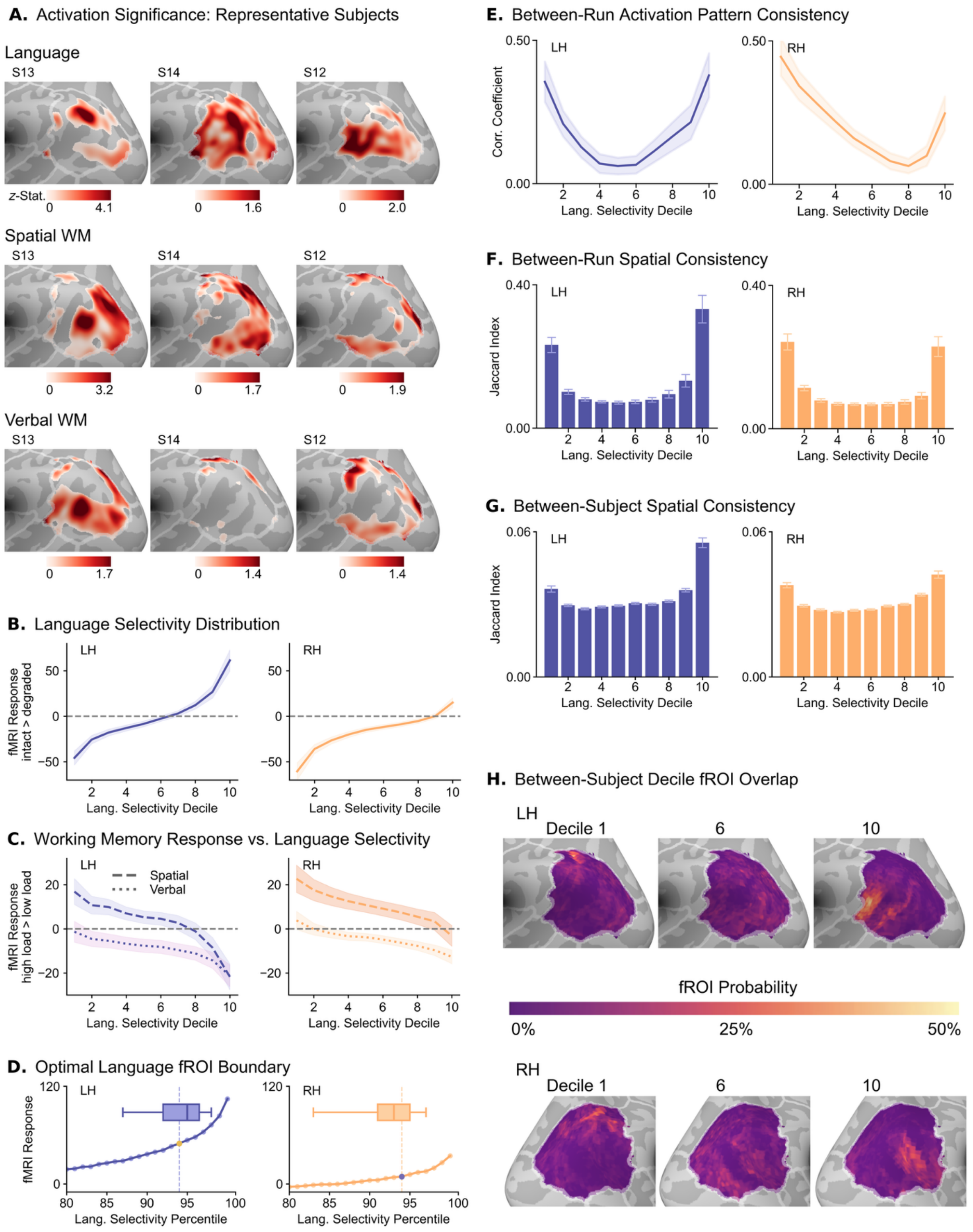


**Supplementary Figure 6**: Inferior Parietal Lobule. **(A – C, E – H)** Same descriptions as in Figure 2. **(D)** The knee point for language selectivity at the group average level (average of individuals' fMRI response values for each percentile; dashed lined indicating knee at: LH: 93%ile, RH: 94%ile) and at the individual level (box plot of individuals' knee values; median: LH: 94%ile, RH: 92%ile).

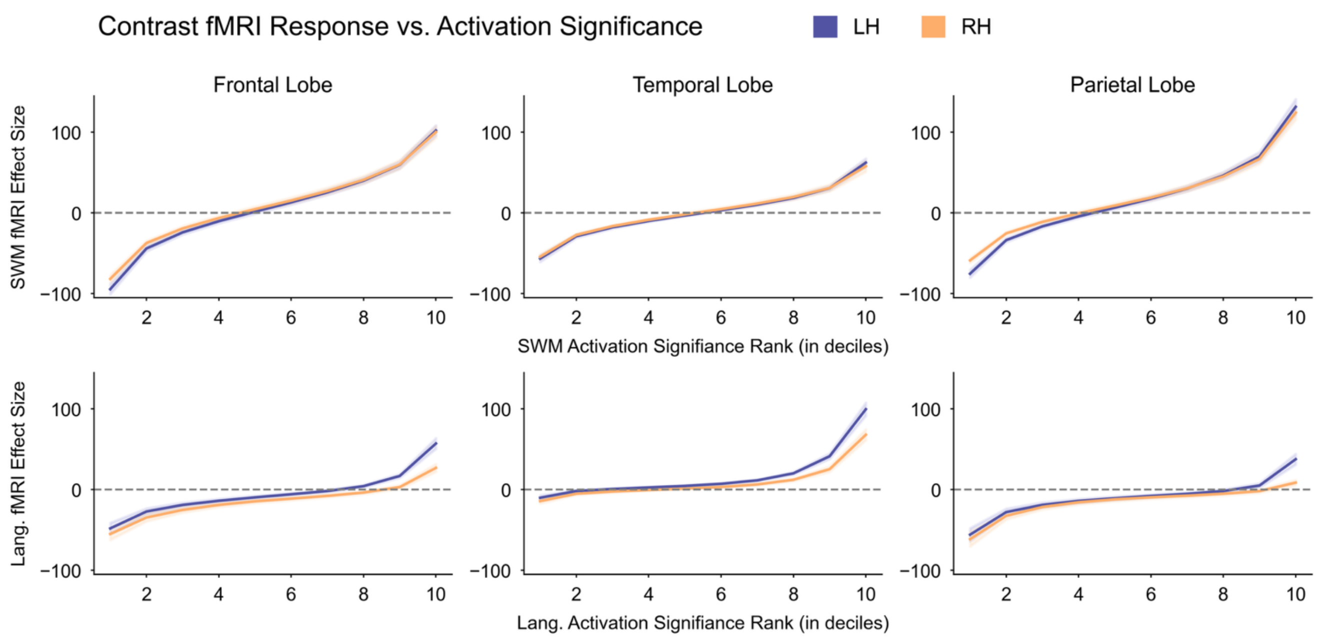


**Supplementary Figure 7**: The fMRI effect size (contrast parameter estimate) averaged within each decile fROI (defined based on the activation significance values) for the Spatial Working Memory (hard > easy) contrast (top row) and the Language Localizer (intact > degraded speech) contrast (bottom row). Group averages of individual selectivity curves are shown for the frontal lobe (left column), temporal lobe (middle column), and parietal lobe (right column), with the error ribbon indicating the standard error of the group average. Note that, in contrast to language, the spatial working memory response distribution is considerably more gradient, particularly across areas of intermediate task response.

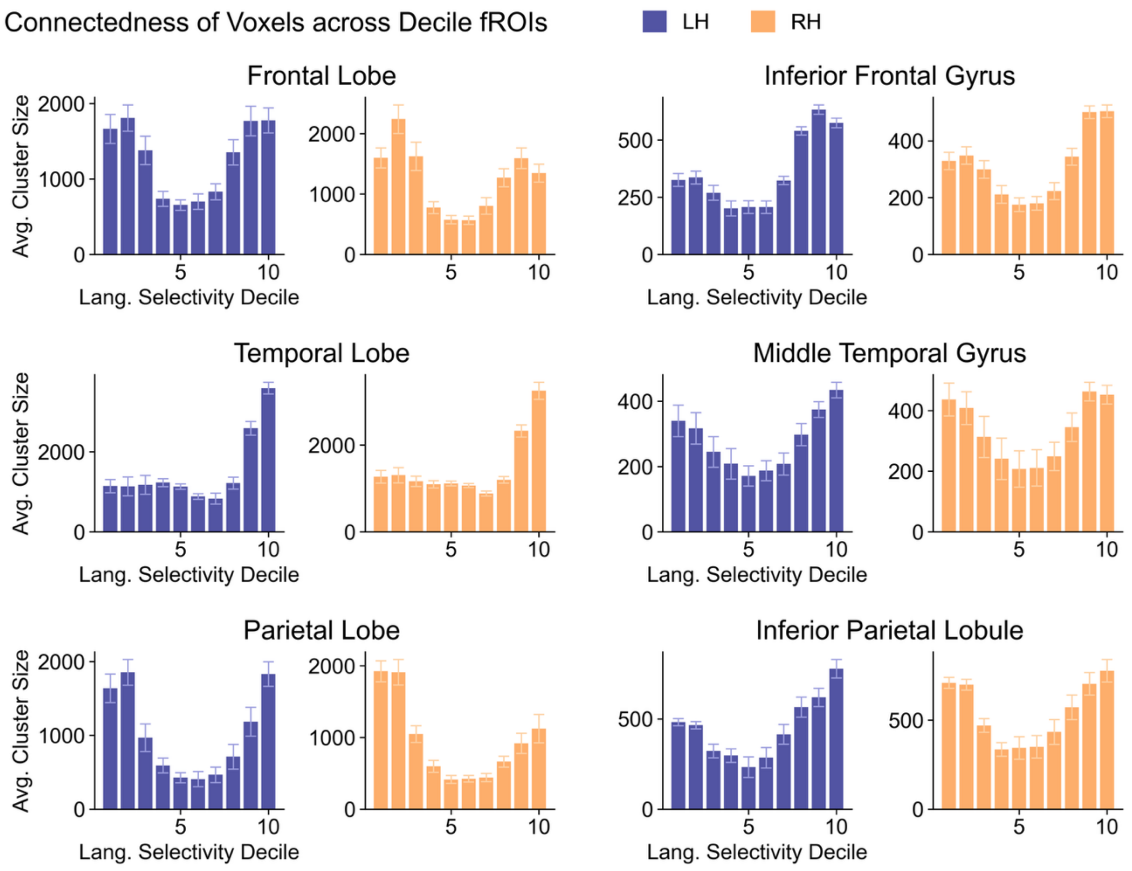


**Supplementary Figure 8**: Within-fROI contiguity by language selectivity decile. The average cluster size (number of adjacent voxels) in each of the 10 language-selectivity decile fROIs is shown for the three lobes and one subregion per lobe. We weighted each cluster size by the proportion of fROI voxels included in that cluster to better reflect the typical connectedness of an fROI voxel.

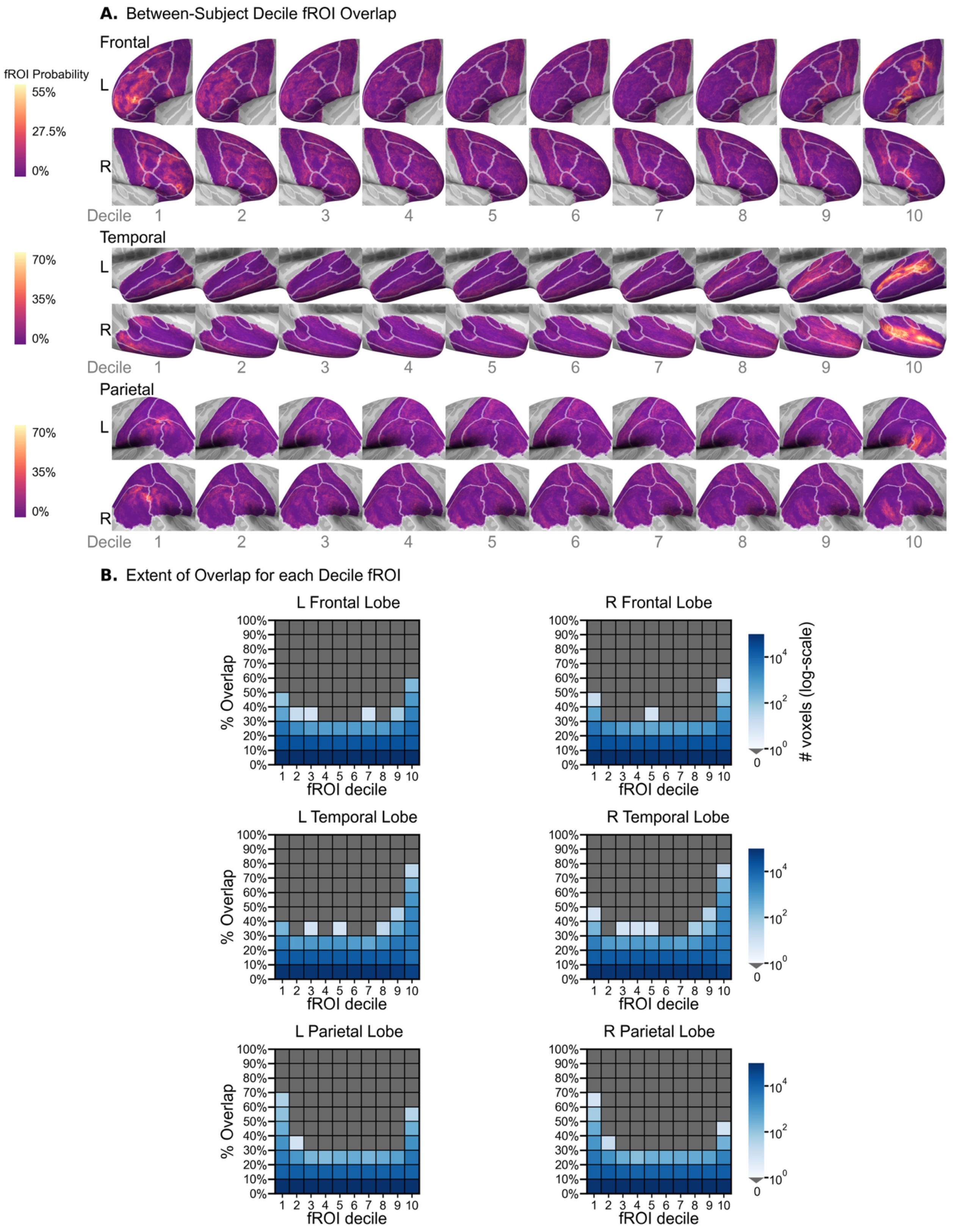


**Supplementary Figure 9**: fROI overlap across subjects by decile. **(A)** Probability maps were created by adding up the binary fROI volumes across individuals and runs, then dividing by the total number volumes. These volumes were then projected to the *fsaverage* surface for visualization. **(B)** Matrices depict the nonzero probability map values (% overlap across individuals and runs) for each of the decile fROIs with the color scale indicating the number of voxels (log-scale) within each interval of % overlap values.

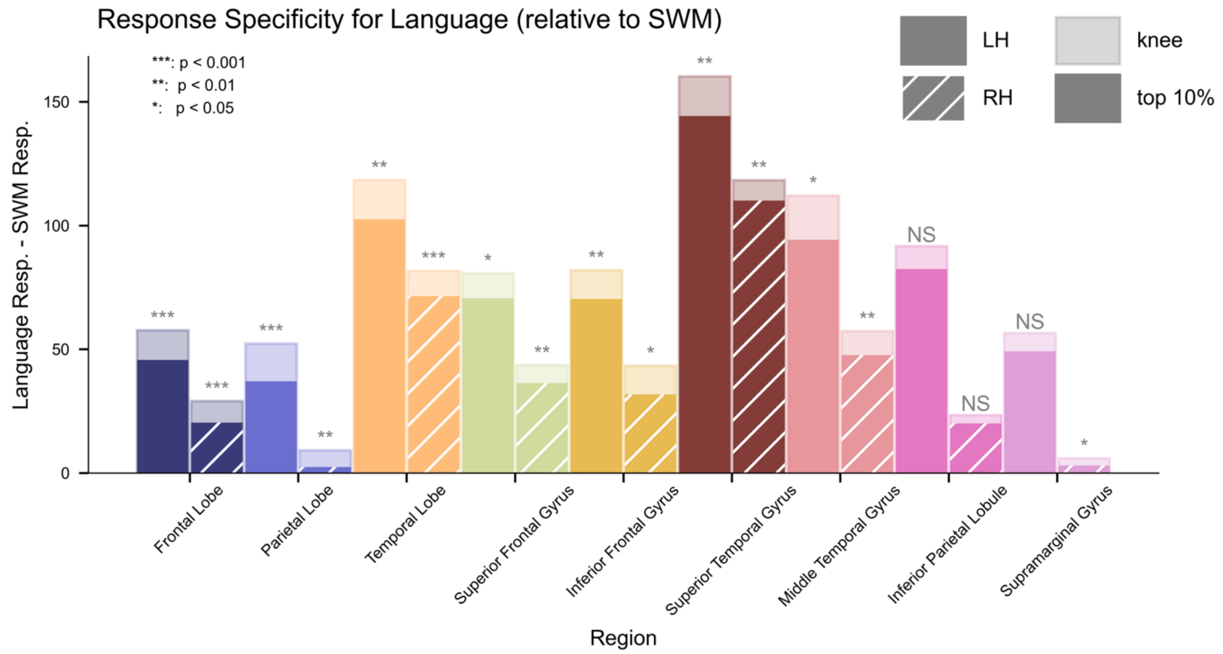


**Supplementary Figure 10**: The difference in fMRI response between the language localizer and the spatial working memory contrasts for the knee fROI and the top 10% fROI. Each significance indicator corresponds to the Bonferroni-corrected *p*-value from a linear mixed effects model assessing how fMRI response difference varies with fROI method (knee or top 10%), with subject as a random effect.

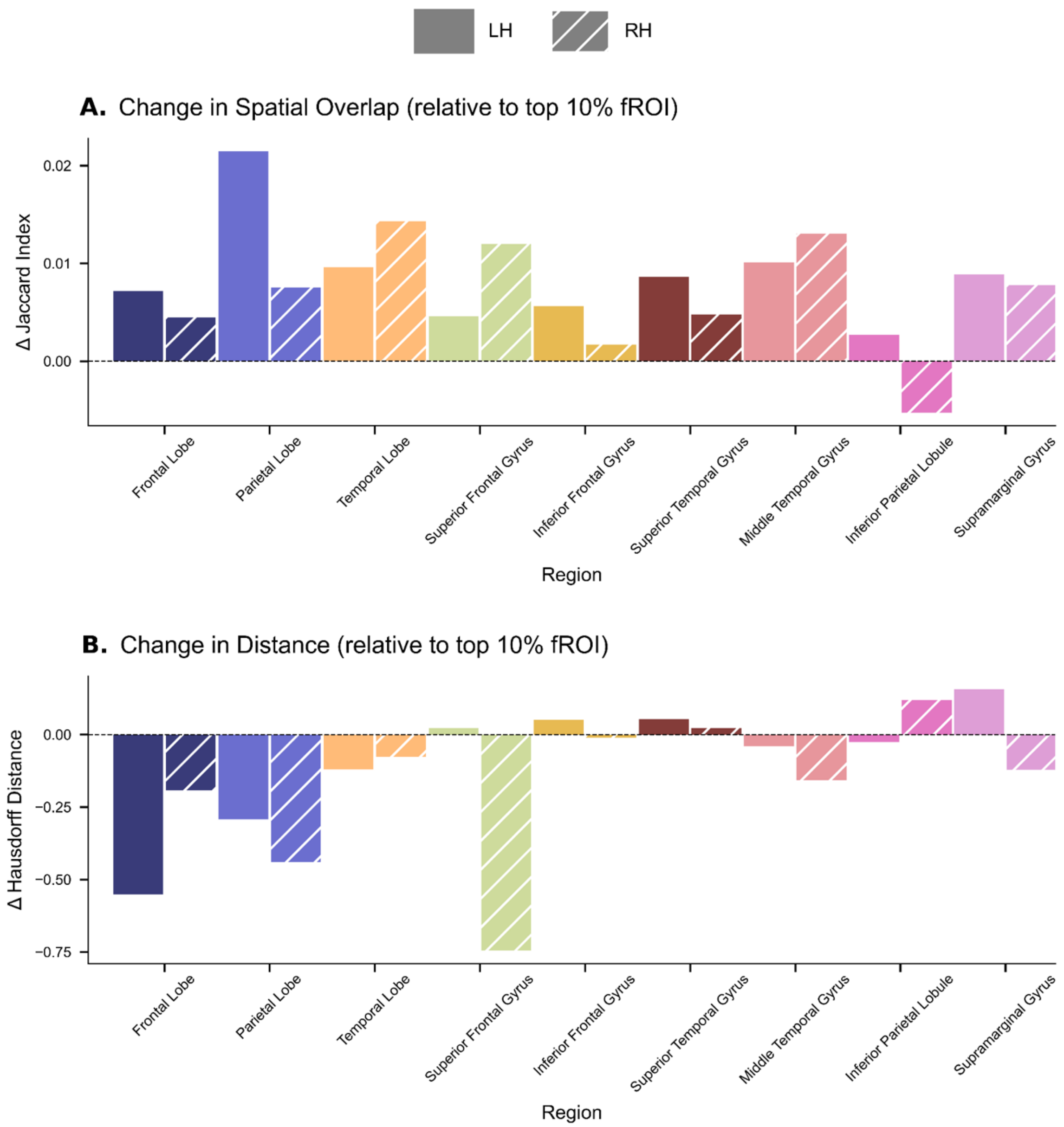


**Supplementary Figure 11**: Comparison of knee fROI to top 10% fROI for test-retest localization. **(A)** The difference in between-run fROI overlap using the knee fROI approach and that of the top 10% approach. A positive change in Jaccard index indicates greater spatial overlap for the knee fROI. **(B)** The difference in between-run fROI Hausdorff distance for the knee fROIs compared to the top 10% fROI. Negative values indicate that the distance between the fROIs from each run decreased.

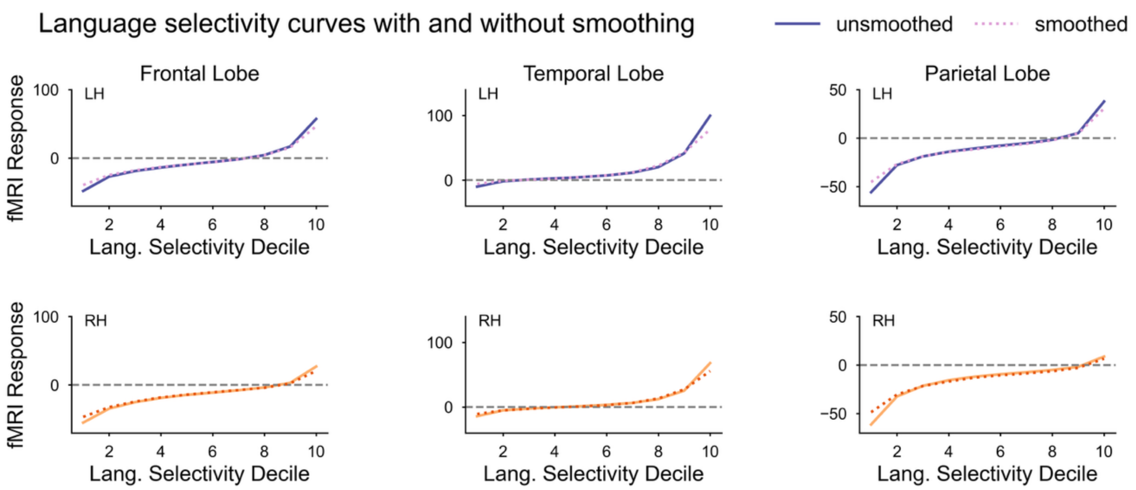


**Supplementary Figure 12**: Language selectivity curves using unsmoothed and smoothed data. The average language localizer fMRI response (intact > degraded) within each decile fROI was calculated from each individual's fMRI data that had either been smoothed (6mm FWHM kernel, FSL's SUSAN) or not smoothed during preprocessing. The decile-specific values were averaged across individuals, and the group average values were plotted.

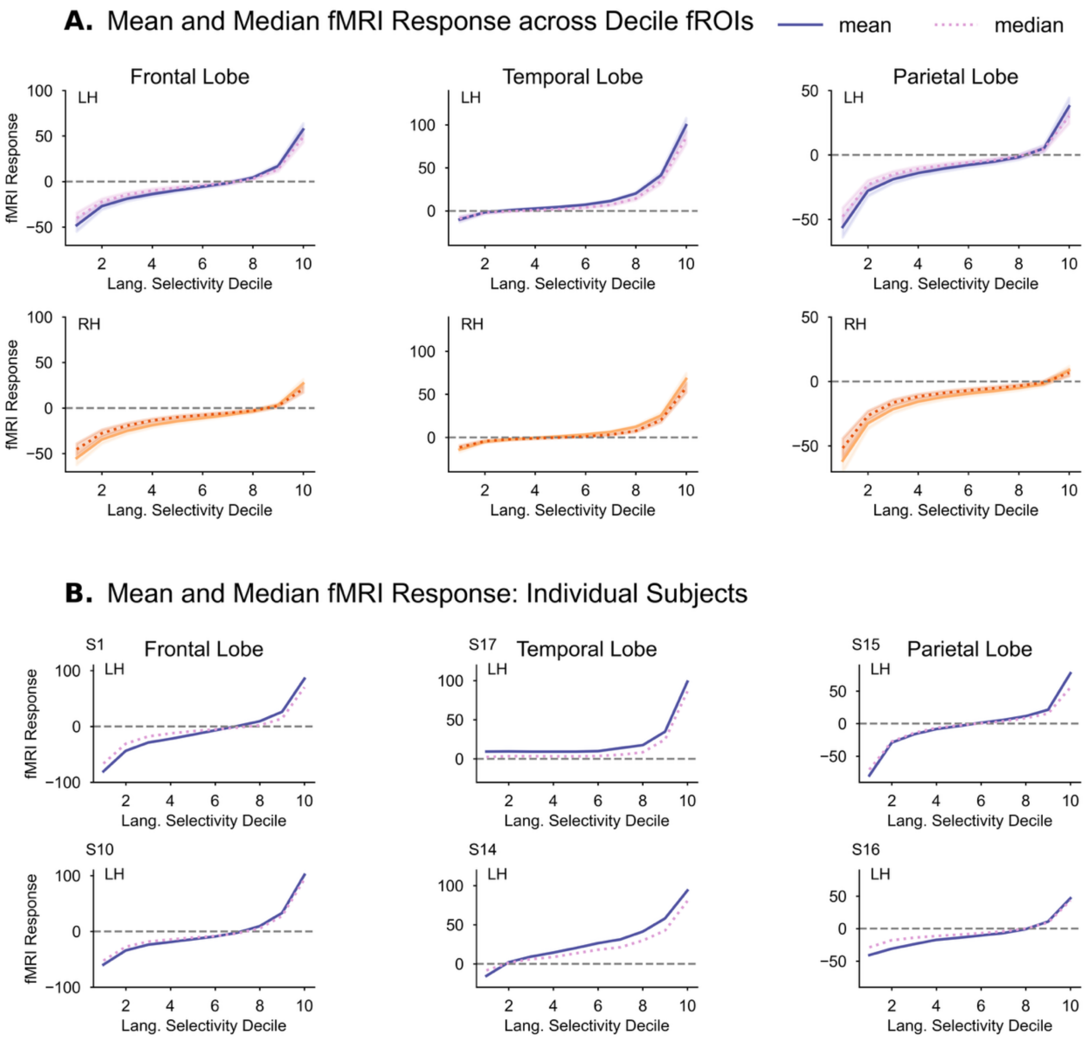


**Supplementary Figure 13**: The mean or median fMRI response value within each language activation-decile fROI. **(A)** Group averages of the decile-specific means or medians. The ribbons reflect the standard error for the group average of the mean values and the group average of the median values. The correspondence between the two curves (median-based, mean-based) shows that the nonlinear shape is not simply attributable to extreme values influencing the means. **(B)** Two example individuals for each left hemisphere region.

**Supplementary Table 1:** Summary of results of the linear mixed effects model for the cubic contrast of language localizer fMRI response against fROI decile. The reported *p*-values have been Bonferroni corrected for the 18 tests (one per region). The relationship (language fMRI response vs. language selectivity decile) was significantly cubic for all regions tested.

| Hemisphere | Region | Estimate ($x^3$) | SE ($x^3$) | *t*-value ($x^3$) | *p*-value ($x^3$) |
|---|---|---|---|---|---|
| LH | Frontal Lobe | 27.713 | 2.849 | 9.726 | $<3.60 \times 10^{-15}$ |
| RH | Frontal Lobe | 19.148 | 2.648 | 7.230 | $3.10 \times 10^{-11}$ |
| LH | Temporal Lobe | 32.606 | 2.311 | 14.110 | $<3.60 \times 10^{-15}$ |
| RH | Temporal Lobe | 24.554 | 1.840 | 13.344 | $< 3.60 \times 10^{-15}$ |
| LH | Parietal Lobe | 27.629 | 3.077 | 8.979 | $< 3.60 \times 10^{-15}$ |
| RH | Parietal Lobe | 17.877 | 2.596 | 6.886 | $2.97 \times 10^{-10}$ |
| LH | IFG | 18.768 | 3.593 | 5.223 | $4.59 \times 10^{-6}$ |
| RH | IFG | 16.969 | 3.216 | 5.275 | $3.49 \times 10^{-6}$ |
| LH | SFG | 23.731 | 3.459 | 6.860 | $3.53 \times 10^{-10}$ |
| RH | SFG | 16.821 | 3.018 | 5.574 | $7.18 \times 10^{-7}$ |
| LH | STG | 20.193 | 3.244 | 6.225 | $1.79 \times 10^{-8}$ |
| RH | STG | 18.393 | 2.611 | 7.045 | $1.06 \times 10^{-10}$ |
| LH | MTG | 25.794 | 2.889 | 8.930 | $< 3.60 \times 10^{-15}$ |
| RH | MTG | 12.660 | 2.125 | 5.958 | $8.44 \times 10^{-8}$ |
| LH | SMG | 26.683 | 3.662 | 7.287 | $2.12 \times 10^{-11}$ |
| RH | SMG | 14.374 | 2.822 | 5.094 | $8.82 \times 10^{-6}$ |
| LH | Inf Parietal | 23.214 | 4.266 | 5.441 | $1.47 \times 10^{-6}$ |
| RH | Inf Parietal | 16.704 | 2.982 | 5.601 | $6.19 \times 10^{-7}$ |

**Supplementary Table 2:** Summary of results of the linear mixed effects model for the quadratic contrast of LangLoc between-run correlation against fROI decile. The reported *p*-values have been Bonferroni corrected for the 18 tests (one per region). The relationship (between-run correlation vs. language selectivity decile) was significantly quadratic for all regions tested.

| **Hemisphere** | **Region** | **Estimate ($x^2$)** | **SE ($x^2$)** | ***t*-value ($x^2$)** | ***p*-value ($x^2$)** |
|---|---|---|---|---|---|
| LH | Frontal Lobe | 0.372 | 0.022 | 16.606 | $< 3.60 \times 10^{-15}$ |
| RH | Frontal Lobe | 0.321 | 0.025 | 14.284 | $< 3.60 \times 10^{-15}$ |
| LH | Temporal Lobe | 0.334 | 0.022 | 15.346 | $< 3.60 \times 10^{-15}$ |
| RH | Temporal Lobe | 0.314 | 0.019 | 16.737 | $< 3.60 \times 10^{-15}$ |
| LH | Parietal Lobe | 0.390 | 0.028 | 14.135 | $< 3.60 \times 10^{-15}$ |
| RH | Parietal Lobe | 0.226 | 0.028 | 8.168 | $4.27 \times 10^{-14}$ |
| LH | IFG | 0.289 | 0.026 | 11.068 | $< 3.60 \times 10^{-15}$ |
| RH | IFG | 0.206 | 0.027 | 7.693 | $1.29 \times 10^{-12}$ |
| LH | SFG | 0.349 | 0.033 | 10.574 | $< 3.60 \times 10^{-15}$ |
| RH | SFG | 0.342 | 0.029 | 11.819 | $< 3.60 \times 10^{-15}$ |
| LH | STG | 0.153 | 0.023 | 6.682 | $1.09 \times 10^{-9}$ |
| RH | STG | 0.146 | 0.022 | 6.590 | $1.94 \times 10^{-9}$ |
| LH | MTG | 0.304 | 0.025 | 11.944 | $< 3.60 \times 10^{-15}$ |
| RH | MTG | 0.228 | 0.024 | 9.549 | $< 3.60 \times 10^{-15}$ |
| LH | SMG | 0.365 | 0.028 | 13.061 | $< 3.60 \times 10^{-15}$ |
| RH | SMG | 0.179 | 0.027 | 6.528 | $2.84 \times 10^{-9}$ |
| LH | Inf Parietal | 0.339 | 0.031 | 10.830 | $< 3.60 \times 10^{-15}$ |
| RH | Inf Parietal | 0.232 | 0.026 | 8.769 | $< 3.60 \times 10^{-15}$ |

**Supplementary Table 3:** Linear mixed effects model results for the within-subject, between-run overlap (Jaccard index) of the 90%ile fROI vs. 80%ile fROI. The reported *p*-values have been Bonferroni corrected for the 18 tests (one per region). The 90%ile fROI had significantly greater between-run overlap than the 80%ile fROI for all regions tested.

| Hemisphere | Region | Estimate | SE | *t*-value | *p*-value |
|---|---|---|---|---|---|
| LH | Frontal Lobe | 0.202 | 0.020 | 9.84 | $3.12 \times 10^{-14}$ |
| RH | Frontal Lobe | 0.135 | 0.017 | 8.13 | $7.50 \times 10^{-11}$ |
| LH | Temporal Lobe | 0.279 | 0.026 | 10.9 | $2.57 \times 10^{-16}$ |
| RH | Temporal Lobe | 0.246 | 0.021 | 11.6 | $1.33 \times 10^{-17}$ |
| LH | Parietal Lobe | 0.179 | 0.021 | 8.57 | $1.03 \times 10^{-11}$ |
| RH | Parietal Lobe | 0.091 | 0.014 | 6.32 | $2.37 \times 10^{-7}$ |
| LH | IFG | 0.197 | 0.026 | 7.62 | $7.62 \times 10^{-10}$ |
| RH | IFG | 0.159 | 0.019 | 8.50 | $1.37 \times 10^{-11}$ |
| LH | SFG | 0.174 | 0.020 | 8.78 | $3.92 \times 10^{-12}$ |
| RH | SFG | 0.149 | 0.019 | 7.74 | $4.46 \times 10^{-10}$ |
| LH | STG | 0.223 | 0.027 | 8.38 | $2.46 \times 10^{-11}$ |
| RH | STG | 0.191 | 0.022 | 8.56 | $1.07 \times 10^{-11}$ |
| LH | MTG | 0.220 | 0.028 | 7.88 | $2.33 \times 10^{-10}$ |
| RH | MTG | 0.155 | 0.019 | 8.19 | $5.80 \times 10^{-11}$ |
| LH | SMG | 0.205 | 0.026 | 7.76 | $4.01 \times 10^{-10}$ |
| RH | SMG | 0.097 | 0.017 | 5.61 | $5.02 \times 10^{-6}$ |
| LH | Inf Parietal | 0.200 | 0.028 | 7.24 | $4.16 \times 10^{-9}$ |
| RH | Inf Parietal | 0.137 | 0.022 | 6.20 | $4.09 \times 10^{-7}$ |

**Supplementary Table 4:** Linear mixed effects model results for between-subject overlap (Jaccard index) of the 90%ile fROI vs. 80%ile fROI. The reported *p*-values have been Bonferroni corrected for the 18 tests (one per region). The 90%ile fROI had significantly greater between-subject overlap than the 80%ile fROI in the temporal lobe only.

| Hemisphere | Region | Estimate | SE | *t*-value | *p*-value |
|---|---|---|---|---|---|
| LH | Frontal Lobe | 0.036 | 0.020 | 1.75 | n.s. |
| RH | Frontal Lobe | 0.016 | 0.017 | 0.94 | n.s. |
| LH | Temporal Lobe | 0.087 | 0.026 | 3.40 | $1.89 \times 10^{-2}$ |
| RH | Temporal Lobe | 0.085 | 0.021 | 4.00 | $2.49 \times 10^{-3}$ |
| LH | Parietal Lobe | 0.027 | 0.021 | 1.28 | n.s. |
| RH | Parietal Lobe | 0.009 | 0.014 | 0.63 | n.s. |
| LH | IFG | 0.017 | 0.026 | 0.66 | n.s. |
| RH | IFG | 0.009 | 0.019 | 0.47 | n.s. |
| LH | SFG | 0.031 | 0.020 | 1.56 | n.s. |
| RH | SFG | 0.019 | 0.019 | 0.99 | n.s. |
| LH | STG | 0.023 | 0.027 | 0.85 | n.s. |
| RH | STG | 0.029 | 0.022 | 1.29 | n.s. |
| LH | MTG | 0.013 | 0.028 | 0.46 | n.s. |
| RH | MTG | 0.017 | 0.019 | 0.90 | n.s. |
| LH | SMG | 0.012 | 0.026 | 0.46 | n.s. |
| RH | SMG | 0.004 | 0.017 | 0.22 | n.s. |
| LH | Inf Parietal | 0.020 | 0.028 | 0.71 | n.s. |
| RH | Inf Parietal | 0.008 | 0.022 | 0.38 | n.s. |

**Supplementary Table 5:** Linear mixed effects model results for the interaction of within-subject vs. between-subject overlap and 80%ile vs. 90%ile overlap. The reported *p*-values have been Bonferroni corrected for the 18 tests (one per region). The increase in overlap for the 90%ile fROI relative to the 80%ile fROI was significantly larger for the within-subject than the between-subject measure for all regions tested.

| **Hemisphere** | **Region** | **Estimate** | **SE** | ***t*-value** | ***p*-value** |
|---|---|---|---|---|---|
| LH | Frontal Lobe | 0.083 | 0.014 | 81.0 | $3.12 \times 10^{-6}$ |
| RH | Frontal Lobe | 0.060 | 0.012 | 81.0 | $4.23 \times 10^{-5}$ |
| LH | Temporal Lobe | 0.096 | 0.018 | 81.0 | $1.68 \times 10^{-5}$ |
| RH | Temporal Lobe | 0.081 | 0.015 | 81.0 | $1.39 \times 10^{-5}$ |
| LH | Parietal Lobe | 0.076 | 0.015 | 81.0 | $3.23 \times 10^{-5}$ |
| RH | Parietal Lobe | 0.041 | 0.010 | 81.0 | $2.30 \times 10^{-3}$ |
| LH | IFG | 0.090 | 0.018 | 81.0 | $8.08 \times 10^{-5}$ |
| RH | IFG | 0.075 | 0.013 | 81.0 | $3.67 \times 10^{-6}$ |
| LH | SFG | 0.071 | 0.014 | 81.0 | $3.92 \times 10^{-5}$ |
| RH | SFG | 0.065 | 0.014 | 81.0 | $1.43 \times 10^{-4}$ |
| LH | STG | 0.100 | 0.019 | 81.0 | $1.61 \times 10^{-5}$ |
| RH | STG | 0.081 | 0.016 | 81.0 | $3.36 \times 10^{-5}$ |
| LH | MTG | 0.104 | 0.020 | 81.0 | $2.17 \times 10^{-5}$ |
| RH | MTG | 0.069 | 0.013 | 81.0 | $3.23 \times 10^{-5}$ |
| LH | SMG | 0.096 | 0.019 | 81.0 | $3.03 \times 10^{-5}$ |
| RH | SMG | 0.047 | 0.012 | 81.0 | $4.84 \times 10^{-3}$ |
| LH | Inf Parietal | 0.090 | 0.020 | 81.0 | $2.60 \times 10^{-4}$ |
| RH | Inf Parietal | 0.064 | 0.016 | 81.0 | $1.66 \times 10^{-3}$ |
